# AMPK activation induces RALDH^high^ tolerogenic dendritic cells through rewiring of glucose and lipid metabolism

**DOI:** 10.1101/2023.07.04.547639

**Authors:** E. C. Brombacher, T. A. Patente, A. J. van der Ham, T. J. A. Moll, F. Otto, F. W. M. Verheijen, E. A. Zaal, A.H. de Ru, R. T. N. Tjokrodirijo, C. R. Berkers, P. A. van Veelen, B. Guigas, B. Everts

## Abstract

It is well known that dendritic cell (DC) activation and function are underpinned by profound changes in cellular metabolism. Several studies indicate that the ability of DCs to promote tolerance is dependent on catabolic metabolism. The AMP-activated kinase (AMPK) is a central nutrient and energy sensor whose activation promotes catabolism while inhibiting ATP-consuming anabolic pathways. Yet the contribution of AMPK activation to DC tolerogenicity remains unknown. Here, we show that AMPK activation renders human monocyte-derived DCs tolerogenic as evidenced by an enhanced ability to drive differentiation of regulatory T cells, a process dependent on increased RALDH activity. This is accompanied by a number of distinct metabolic changes, in particular increased breakdown of glycerophospholipids, enhanced mitochondrial fission-dependent fatty acid oxidation, and upregulated glucose catabolism. This metabolic rewiring is functionally important as we found interference with these metabolic processes to reduce to various degrees AMPK-induced RALDH activity as well as the tolerogenic capacity of moDCs. Altogether, our findings reveal a key role for AMPK signaling in shaping DC tolerogenicity, and suggest that AMPK may serve as new target to direct DC-driven immune responses in therapeutic settings.

## Introduction

Dendritic cells (DCs) are specialized antigen-presenting cells with the ability to prime and polarize T cell responses. As such, DCs play a central role in the establishment of protective adaptive immunity following infections and after vaccination. In addition, their ability to prime regulatory T cells (Tregs) and induce anergy to host self-antigens make them critical regulators of tolerance^1^. Hence, promoting tolerogenic DCs (tolDCs) could be an interesting therapeutic strategy for inflammatory diseases. Although several clinical studies have already confirmed the safety of tolDC-based therapies, the clinical benefits are currently limited, in part due to impaired long-lasting maintenance of the tolerogenic state^2–4^. Novel insights into the processes that drive and shape DC tolerogenicity are therefore instrumental for devising new strategies to increase efficacy of tolDC-based therapies.

DCs can be exposed to a number of different metabolic micro-environments ranging from a nutrient-deprived tumor micro-environment to nutrient-rich tissues in obese individuals. These metabolic cues and nutrient availability are expected to have a major impact on DC function and the subsequent immune response, as they impinge on intracellular metabolic pathway activity that play a central role in shaping DC immunogenicity^5^. Acquisition of an immunogenic DC phenotype is commonly associated with anabolic metabolism and underpinned by enhanced glycolysis and fatty acid synthesis^6,7^, while tolDCs are functionally supported by catabolism-centered metabolism, characterized by enhanced glycolysis in combination with elevated oxidative phosphorylation and fatty acid oxidation (FAO)^8–10^.

Nutrient-sensing pathways connect environmental cues to changes in cellular bioenergetics and are therefore highly important for DC function. For example, DC activation is facilitated by upregulated glycolysis, which is supported by mechanistic target of rapamycin (mTOR) and counteracted by AMP-activated protein kinase (AMPK)^7^, two nutrient sensing enzymes that are activated in nutrient-sufficient and -poor conditions, respectively^11,12^. While the role of mTOR in DCs has been extensively studied, less is known about the role of AMPK in DC function^12^. AMPK is a central regulator of cellular metabolism, well known for its role in regulating energy homeostasis during nutrient stress. Canonical AMPK activation involves a high AMP/ADP ratio that triggers Liver Kinase B1 (LKB1) to activate AMPK via phosphorylation. AMPK activation inhibits fatty acid synthesis while enhancing FAO, by phosphorylation of acetyl-CoA carboxylase (ACC) isoforms ACC1 and ACC2, respectively. Catabolic metabolism is further promoted by induction of autophagy and downregulating of protein synthesis through inhibition of mTORC1. AMPK also plays an important role in mitochondrial homeostasis by regulating the removal of damaged mitochondria through mitophagy and by stimulating mitochondrial biogenesis^11,13^.

There is some evidence that suggests a role for AMPK signaling in DCs in dampening an inflammatory immune response. AMPK activation in bone-marrow derived DCs (BMDCs) suppressed IL-12p40 secretion and CD86 expression^7^, while AMPK deficiency in BMDCs promotes secretion of pro-inflammatory cytokines and IFN-γ and IL-17 secretion by T cells^14^. Additionally, AMPK deficiency in CD11c-expressing cells worsened lung injury following hookworm infection through increased IL- 12/23p40 production by pulmonary CD11c^+^ cells^15^. Interestingly, we have recently found that the tolerogenic effects of retinoic acid (RA) on DCs are AMPK dependent, suggesting that AMPK may not only dampen DC immunogenicity, but can also actively contribute to the tolerogenic capacity of DCs^16^. Yet, whether AMPK activation itself is sufficient to induce DC tolerogenicity and immune tolerance and what the underlying mechanisms are remains unknown.

In the present study, we find that drug-induced AMPK activation in DCs is sufficient to promote their tolerogenicity, as evidenced by an enhanced ability to prime functional regulatory T cells through induction of retinaldehyde dehydrogenase (RALDH) activity. We further show that, mechanistically, AMPK activation boosts glycerophospholipid degradation, mitochondrial-fission induced FAO, and glucose catabolism. Interference with these metabolic alterations partly negate the tolerogenic effects of AMPK activation. Together this provides support for an important role of AMPK in rendering DCs tolerogenic through rewiring of lipid and glucose metabolism and may identify AMPK as a potential target for improving tolDC-based immunotherapies.

## Results

### AMPK activation renders DCs tolerogenic through promotion of RALDH activity

To study the effect AMPK activation on DC function, we treated human monocyte-derived DCs (moDCs) with 991, a direct small-molecule AMPK activator^17^. AMPK activation was rapidly induced by 991, as assessed by phosphorylation on Ser79 of its downstream target Acetyl-CoA Carboxylase (ACC)^18^, and remained stable over time (Supp. Fig. 1A). To investigate the effects of AMPK activation on DC immunogenicity, immature moDCs (iDCs) were treated overnight with 991 before LPS stimulation (Fig. 1A). 24 hours later both iDCs and LPS-treated mature DCs (mDCs) showed increased AMPK activation (Fig. 1B), while simultaneously reducing mTORC1 activity, as assessed by phosphorylation of S6, a downstream target of mTORC1 (Supp. Fig. 1B). Of note, 991 treatment did not affect cell viability (Supp. Fig. 1C). 991 stimulation diminished LPS-induced activation of DCs when compared to DMSO-treated cells, as evidenced by reduction in several activation markers, as well as IL-10 and IL12p40 secretion after co-culture with a CD40L expressing cell line to mimic T cell interactions (Fig. 1C, D and Supp. Fig. 1D,E). Conversely, expression of CD103 was increased (Fig. 1C), a marker also upregulated on RA-induced tolerogenic DCs which are known to promote Treg differentiation through retinaldehyde dehydrogenase (RALDH)-driven RA production^19,20^. Interestingly, 991 treatment also promoted RALDH activity in mDCs (Fig. 1E).

**Figure 1.**
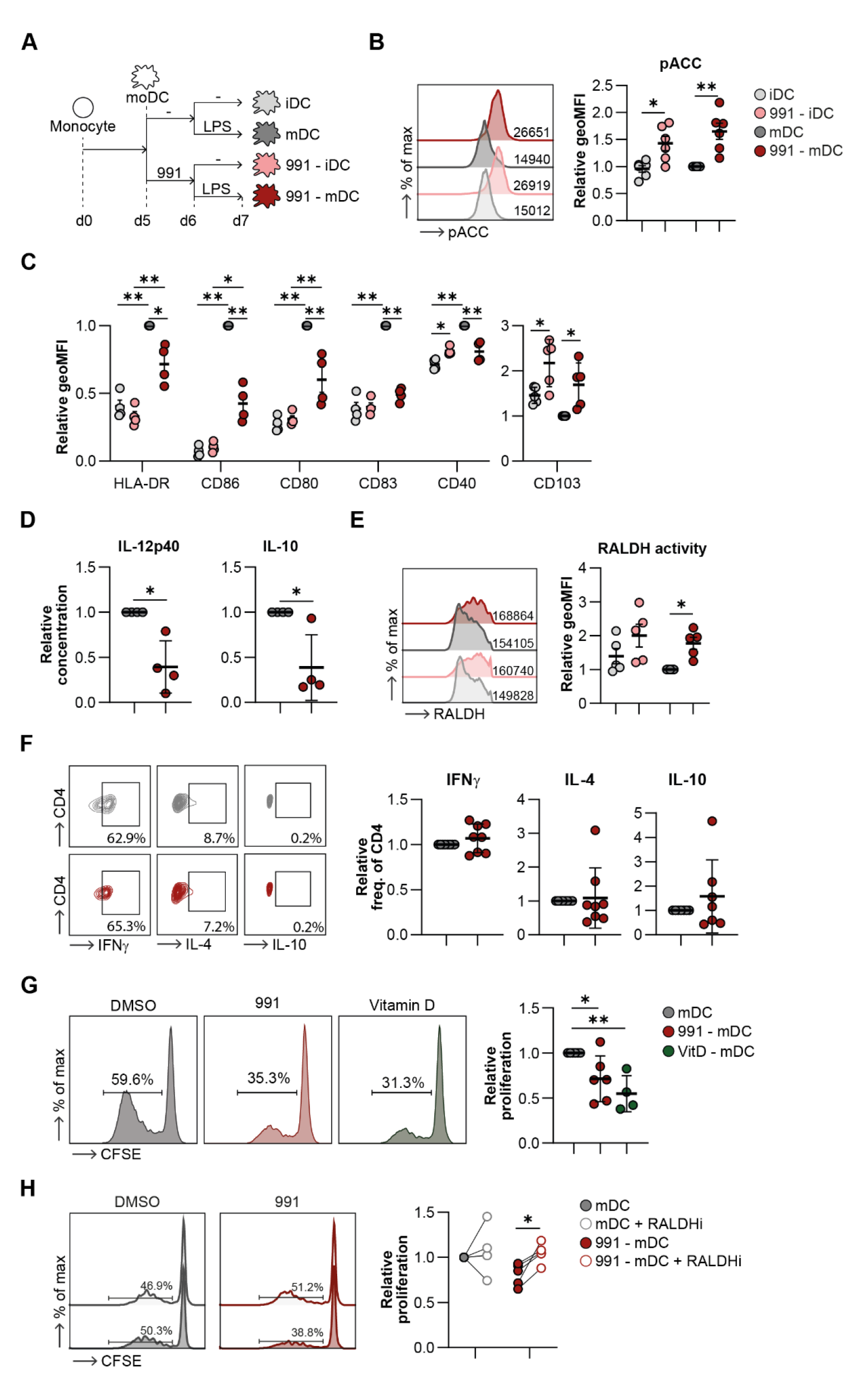
AMPK activation induces tolerogenic DCs via promotion of RALDH activity. **A:** Schematic overview of experimental set-up. Monocytes were isolated from PBMCs and differentiated in moDCs in the presence of GM-CSF + IL-4. On day 5 cells were treated with either DMSO or 991. On day 6, 100 ng/mL of LPS was added if indicated and 24 hours later cells were harvested for functional assays. **B:** Representative histogram and normalized quantification of phosphorylation of ACC (Ser79). **C:** Normalized expression of indicated markers by moDCs. **D:** Relative concentration of IL-10 and IL12p40 in the supernatants of mDCs after a 24 hour co-culture with CD40L-expressing J558 cells. **E:** Representative histogram and normalized quantification of RALDH activity. **F:** Representative plots and normalized percentages of CD4^+^ T cells of intracellular cytokines after restimulation with PMA/ionomycin in the presence of Brefeldin A. **G, H:** Representative histograms and normalized percentage of proliferation of bystander T cells after co-culture with irradiated T cells primed with **(G)** 991/DMSO-treated mDCs and **(H)** in presence or absence of RALDH inhibitor (RALDHi) bisdiamine during both mDC maturation and mDC-T cell co-culture. Results are expressed as means ± SEM. Datapoints represent independent experiments with different donors. Statistical analyses were performed using paired t-tests **(D, F)**, one-way Anova with Sidak post-hoc test **(G)** or two-way Anova with Tukey post-hoc test **(B, C, E, H)**. *p < 0.05, **p < 0.01.

Next, the effect of AMPK activation on T cell priming by mDCs was evaluated. A co-culture with allogenic naïve CD4^+^ T cells showed that the ability of mDCs to promote IFN-γ, IL-4, and IL-10 secretion by T cells was not modulated by 991 (Fig. 1F and Supp. Fig. 1F). However, the capacity of these T cells to reduce proliferation of bystander T cells was enhanced compared to T cells primed with DMSO-treated mDCs (Fig. 1G), indicating that AMPK activation renders DCs tolerogenic as determined by their ability to prime functional regulatory T cells. A similar trend towards induction of Tregs was observed when DCs were treated with AICA riboside (AICAR), another AMPK activator (Supp. Fig. 1G,H). Of note, further phenotypic characterization of Tregs primed by 991-stimulated mDCs did not reveal enrichment of any of the main previously described subsets of induced Tregs, including CD127^-^CD25^+^FoxP3^+^CTLA4^+^ Tregs (induced Tregs (iTregs))^21^, ICOS^+^ Tregs^22^, TIGIT^+^ Tregs^23^, or Lag3^+^CD49b^+^ Treg1 cells^24^ (Supp. Fig. 1I, J), suggesting that AMPK-activated DCs drive induction of a non-canonical Treg population^21–24^.

We next questioned whether the tolerogenic effects of AMPK activation were dependent on increased RALDH activity. mDCs were therefore treated with RALDH2 inhibitor bisdiamide during 991 treatment and subsequent LPS stimulation (Supp. Fig. 1K). RALDH inhibition prevented 991-induced suppression in expression of HLA-DR, CD86, CD83 and CD80 to various degrees (Supp. Fig. 1L). IL-10 and IL-12p40 secretion was further decreased upon RALDH inhibition compared to control groups (Supp. Fig. 1M). The suppressive capacity of T cells primed with 991-treated DCs was not affected when RALDH was inhibited during DC culture alone (Supp. Fig. 1N), but strikingly, when bisdiamide was also added during T cell co-culture, the ability of 991-stimulated mDCs to prime functional Tregs was lost. Since T cells did not show detectable RALDH activity (Supp. Fig. 1K), this indicates that RALDH activity in 991-stimulated mDCs during DC-T cell coculture is required for these DCs to exert their tolerogenic effect (Fig. 1H). Altogether, AMPK activation in DCs induces tolerogenicity and promotes priming of non-canonical functional regulatory T cells through induction of RALDH activity.

### AMPK activation drives catabolic glycerophospholipid metabolism

To gain insight into how AMPK activation induces DC tolerogenicity we performed untargeted metabolomics and proteomics on DMSO-and 991-treated iDCs and mDCs. LPS treatment had clear effects on both the metabolome and the proteome of DCs. Strikingly, LPS stimulation did not lead to any of these changes in 991-treated cells (Supp. Fig. 2A,B), supporting our earlier observations that AMPK activation counteracts LPS-induced DC maturation. 991 treatment promoted increases in several proteins and metabolites irrespective of LPS stimulation, while the downregulated hits upon 991 treatment mainly consisted of proteins and metabolites whose accumulation were induced by LPS (Supp. Fig. 2C-E). Pathway enrichment analysis on the proteome revealed that pathways enriched upon 991 stimulation in LPS-treated mDCs were all associated with (catabolic) lipid metabolism (Fig. 2A), in particular glycero(phospho)lipid catabolism. Metabolomic data also indicated enhanced glycerolipid metabolism, as well as upregulation of the pentose phosphate pathway (Fig. 2B). Integration of the proteomic and metabolomic data sets using web-service ‘genes and metabolites’ (GAM)^25^, created a metabolic network that additionally highlighted changes in amino acid metabolism and TCA cycle metabolites, and further affirmed significant changes in lipid metabolism, including glycero(phospho)lipid metabolism (Fig. 2C). Additionally, most 991-specific metabolite and protein changes in glycero(phospho)lipid metabolism occurred independently of DC maturation (Fig. 2D,E). The proteomic/metabolic network connected an accumulation of glycerol-3-phosphate and a decrease in membrane-lipid phosphatidylcholine (Fig. 2F) to upregulation of two proteins involved in breakdown of glycerophospholipids, namely Patatin-like phospholipase domain-containing protein 6 (PNPLA6) and Glycerophosphocholine Phosphodiesterase 1 (GPCPD1) (Fig. 2G). PNPLA6, also known as Neuropathy Target Esterase (NTE), is a phospholipase that hydrolyzes phosphatidylcholine (PC), thereby releasing fatty acids and glycerophosphocholine^26^. GPCPD1, also known as GDE5, hydrolyzes glycerophosphocholine (GroPCho) to glycerol-3-phosphate (G3P) and choline^27^. Phosphatidylethanolamine (PE), another membrane lipid of which the abundance also decreased upon 991 treatment (Fig. 2F), can also be hydrolyzed to G3P and ethanolamine via PNPLA6 and GPCPD1, albeit with a lower affinity^26,27^. We aimed to determine whether breakdown of PC and PE via PNPLA6 and GPCPD1 contributes to AMPK-induced tolerogenicity and hence, we silenced expression of both *PNPLA6* and *GPCPD1* in iDCs, before adding 991 and LPS, with siRNA. qPCR analysis revealed that 991 induced *PNPLA6* and *GPCPD1* protein expression was mirrored at the transcriptional level (Fig. 2H). Gene silencing strongly reduced expression of both genes, although this was less pronounced in 991-treated cells (Fig. 2H). Expression of activation markers and IL-12p40 and IL-10 secretion was not affected by silencing of *PNPLA6* and *GPCPD1* (Supp. Fig. 2G), while RALDH activity in silenced 991- stimulated mDCs showed a reduction in most donors (Fig. 2I). Correspondingly, 991-driven induction of functional Tregs by mDCs was partly lost after silencing of *PNPLA6* and *GPCPD1* (Fig. 2J). Altogether these data suggest that AMPK activation in mDCs promotes catabolic lipid metabolism and that breakdown of glycerophospholipids, mediated by PNPLA6 and GPCPD1, are contributing to AMPK-induced tolerogenicity.

**Figure 2:**
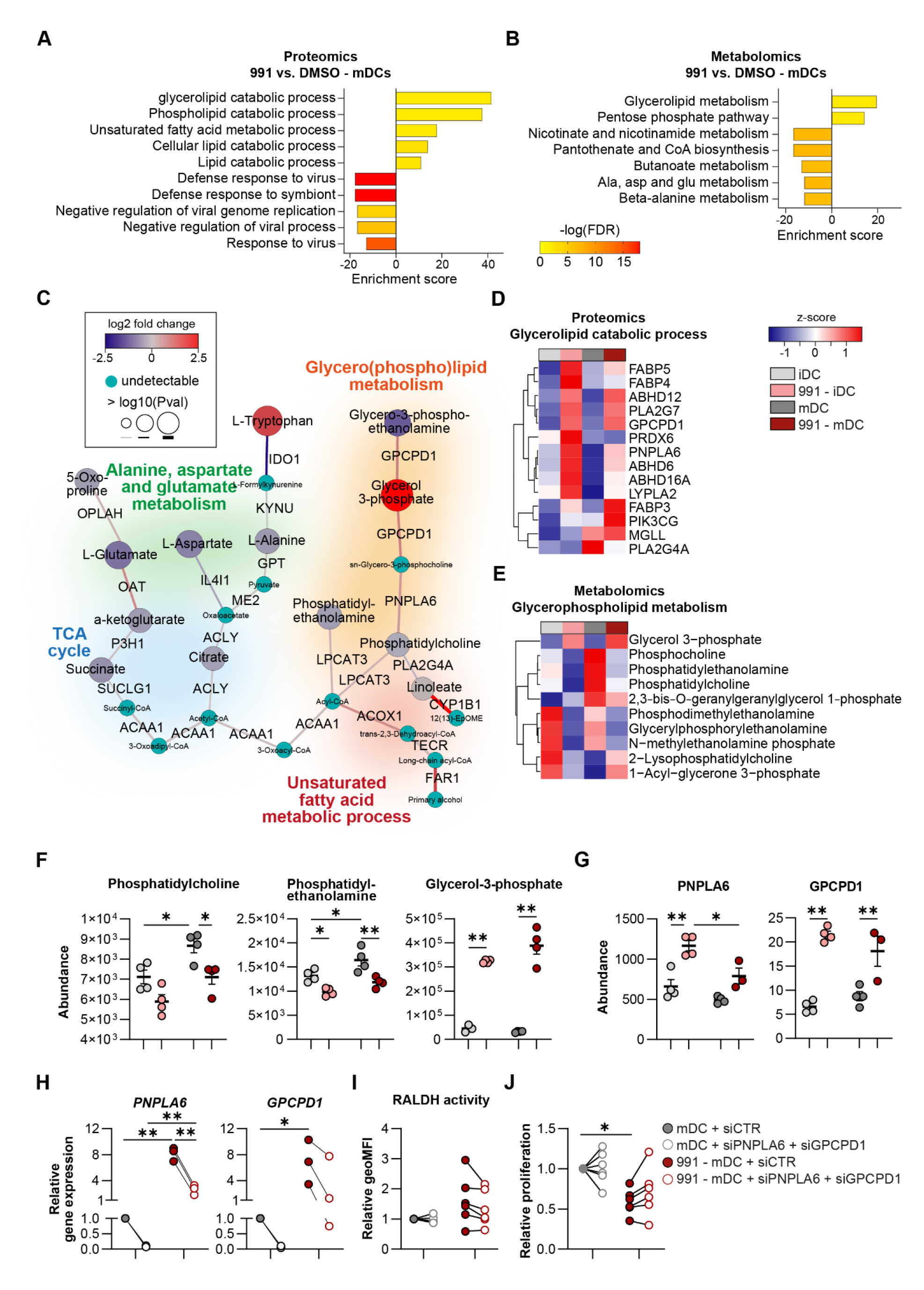
AMPK activation drives glycerophospholipid breakdown. Proteomics and metabolomics were performed on DMSO/991-treated iDCs and mDCs. **A:** Gene ontology (GO) pathway enrichment of significantly up-and downregulated proteins in AMPK-activated mDCs. **B:** KEGG enrichment analysis of significantly up-and downregulated metabolites in AMPK-activated mDCs. **C:** network analysis of integrated proteomics and metabolomics data from mDCs. **D, E:** Heatmap depicting relative expression of **(D)** proteins and **(E)** metabolites involved in glycerophospholipid metabolism. **F, G:** Dotplots showing the abundance as determined by metabolomics or proteomics of **(F)** metabolites and **(G)** proteins involved in glycerophospholipid metabolism that are strongly affected by AMPK activation. **H:** Relative gene expression of *PNPLA6* and *GPCPD1* 72 hours after electroporation with control siRNA (siCTR), or siRNA targeting *PNPLA6* or *GPCPD1*. **I:** normalized quantification of RALDH activity. **J:** Normalized percentage of proliferation of bystander T cells after a T cell suppression assay. Results are expressed as means ± SEM. Datapoints represent independent experiments with different donors. Statistical analyses were performed using paired t-tests **(H)** or two-way Anova with Tukey post-hoc test **(F, G, I, J)**. *p < 0.05, **p < 0.01.

### Fatty acid oxidation is important for AMPK-induced tolerogenicity

As our data indicate enhanced lipid catabolism upon AMPK activation and given that AMPK is known to promote mitochondrial FAO, we further investigated the effects of 991 treatment on mitochondrial metabolism and FAO. Basal mitochondrial respiration, ATP production, and spare respiratory capacity were not affected by AMPK activation (Fig. 3A,B, Supp. Fig. 3A) and neither were mitochondrial mass or membrane potential (Supp. Fig. 3B). Interestingly, 991 treatment increased the protein expression ratio between complex II and complex I of the electron transport chain. This is a metabolic adaptation commonly associated with increased FAO, as it allows for more efficient oxidation of complex II substrate FADH_2_ that is generated during FAO. Hence, this AMPK-activation driven change in ratio may reflect increased FAO rates (Fig. 3C). Indeed, [U-^13^C]-palmitate tracing, revealed a trend towards increased uptake and oxidation of palmitate, marked by higher abundance of labeled Palmitoyl-L-Carnitine and the presence of palmitate-derived carbons in acetyl-CoA, the final product of FAO (Fig. 3D-G). Although the total levels of acetyl-CoA and palmitate-derived acetyl-CoA (M+2) were higher after 991 treatment, the contribution of palmitate to acetyl-CoA was low in these cells, at around 3 percent. AMPK activation did not change carnitine palmitoyltransferase-1A (CPT1A) expression (Supp. Fig. 3C), suggesting that enhanced FAO is driven by higher CPT1A activity and/or by increased free FA availability. In support of the latter, we found that AMPK activation promoted long-chain fatty acid uptake, while lipid droplet accumulation was unaffected (Fig. 3H). Additionally, release of endogenous FAs from phospholipids upon AMPK activation, as shown in Fig. 2, may contribute to increasing intracellular fatty acid levels to fuel enhanced FAO.

**Figure 3:**
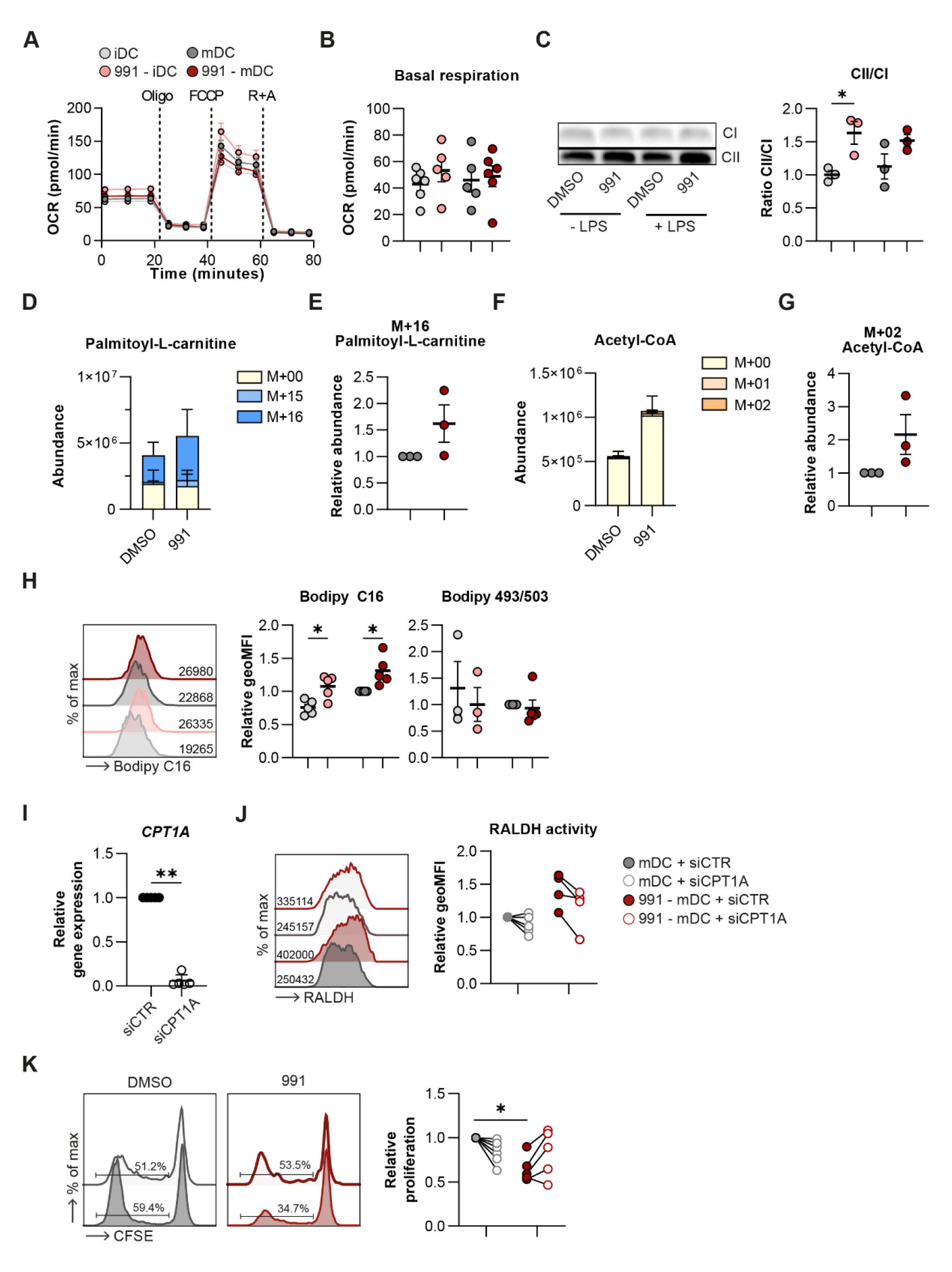
AMPK-induced tolerogenicity is driven by FAO. **A, B:** Real-time oxygen consumption rate (OCR) as measured by Seahorse extracellular flux analysis and **(B)** quantification of basal respiration rates. **C:** Western Blot of complex I (CI) and complex II (CII) of the electron transport chain and quantification of the ratio between CII and CI. **D-G:** [U-^13^C]- palmitate tracing was performed in DMSO/991-treated mDCs. Figures show **(D)** total abundance of palmitoyl-L-carnitine and **(E)** abundance of the labeled M+16 fraction of palmitoyl-L-carnitine normalized per donor and **(F)** total abundance of acetyl-CoA and **(G)** abundance of the labeled M+02 fraction of acetyl-CoA normalized per donor. **H:** representative histogram of Bodipy C16 uptake and normalized quantification of Bodipy C16 and Bodipy 495/503. **I:** Relative gene expression of CPT1A in mDCs 72 hours after electroporation with control siRNA (siCTR) or siRNA targeting CPT1A (siCPT1A). **J:** Representative histogram and normalized quantification of RALDH activity. **K:** Representative histograms and normalized percentage of proliferation of bystander T cells after a T cell suppression assay. Results are expressed as means ± SEM **(B-D, E, G-I)** or means ± SD **(E+G)**. Datapoints represent independent experiments with different donors. Statistical analyses were performed using paired t-tests **(E, G, I)** or two-way Anova with Tukey post-hoc test **(B, C, H, J, K)**. *p < 0.05, **p < 0.01.

To study whether FAO is important for the AMPK-induced Treg-priming capacity of mDCs we blocked mitochondrial import of long-chain fatty acids by silencing *CPT1A* (Fig. 3I). *CPT1A* knockdown did not change uptake of Bodipy C16 and neutral lipid accumulation (Bodipy 493/503), nor did it restore the AMPK-induced inhibition of activation markers and IL-12p40 and IL-10 secretion (Supp. Fig. 3D-F). However, silencing *CPT1A* decreased the AMPK-driven increase in RALDH activity in 991- stimulated mDCs, as well as the suppressive capacity of T cells primed by these DCs (Fig. 3I-K). Taken together, these data show that AMPK activation increases FAO, which promotes RALDH activity and thereby the tolerogenic capacities of moDCs.

### AMPK activation drives fatty acid oxidation through mitochondrial fission

AMPK-induced FAO has been shown to be supported by changes in the mitochondrial fission/fusion balance^28^. We therefore evaluated mitochondrial morphology and questioned whether fission and fusion play a role in the AMPK-driven metabolic and immunological changes. We observed that in both iDCs and mDCs AMPK activation induced fragmentation of mitochondria, indicative of increased fission (Fig. 4A-B). Fusion can be promoted by AMPK through blocking recruitment of DRP1 to the mitochondria via phosphorylation DRP1 (S637)^29^, while phosphorylation of mitochondrial fission factor (MFF) (S172), the receptor for DRP1, promotes fission^30^. 991 treatment did not induce phosphorylation of DRP1 (S637) and counteracted the LPS-induced increase of DRP1 phosphorylation, while it did promote phosphorylation of MFF (S172/S146) in mDCs (Fig. 4C). Furthermore, expression of *FIS1*, a gene involved in mitochondrial fission, increased upon AMPK activation (Supp. Fig. 4A), together providing further support for induction of mitochondrial fission following AMPK activation in moDCs.

**Figure 4:**
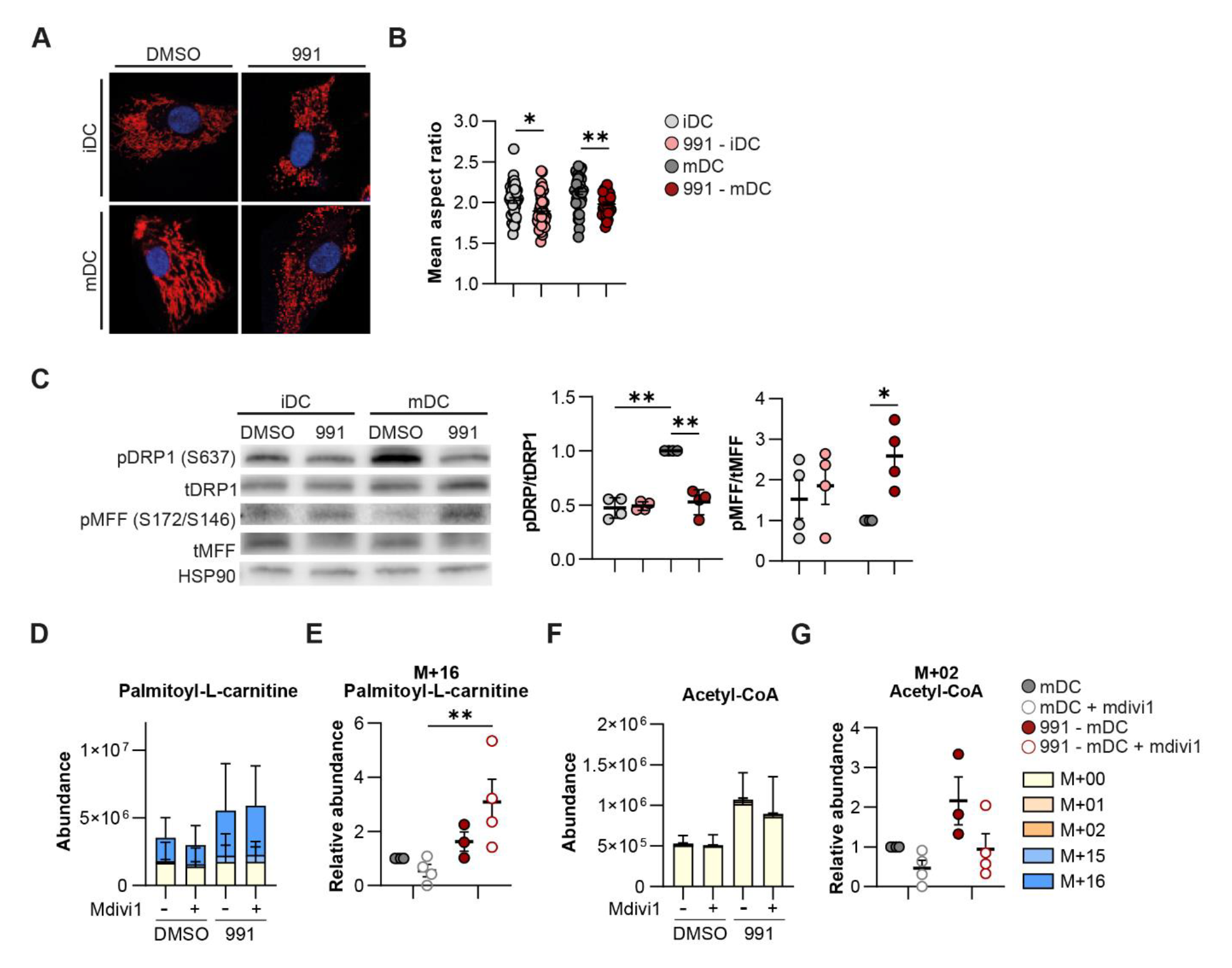
Mitochondrial fission induced by AMPK activation promotes FAO. **A:** Confocal images of DMSO/991-treated iDCs and mDCs. Blue: nucleus. Red: mitochondria. **B:** Quantification of the average aspect ratio of mitochondria per cell. **C:** Western Blot of phosphorylated DRP1 (pDRP1), total DRP1 (tDRP1), phosphorylated MFF (pMFF), total MFF (tMFF), and HSP90 and the ratio for pDRP1/tDRP1 and pMFF/tMFF, normalized for HSP90 and relative to DMSO-treated mDCs. **D-G:** [U-^13^C]-palmitate tracing was performed in DMSO/991-treated mDCs, with or without mdivi1. **(D)** Total abundance of palmitoyl-L-carnitine and **(E)** abundance of the labeled M+16 fraction of palmitoyl-L-carnitine normalized per donor and **(F)** total abundance of acetyl-CoA and **(G)** abundance of the labeled M+02 fraction of acetyl-CoA normalized per donor.Results are expressed as means ± SEM **(B, C, E, G)** or means ± SD **(D,F)**. Datapoints represent independent experiments with different donors. Statistical analyses were performed using two-way Anova with Tukey post-hoc **test (B, C, E, G)**. *p < 0.05, **p < 0.01.

Next, we addressed whether AMPK-driven mitochondrial fission is required for FAO by inhibiting mitochondrial fission using mdivi1, an inhibitor of DRP1 ^31^. Of note, mdivi1 was also shown to inhibit complex I of the electron transport chain, but at the tested concentration in our model, mdivi1 solely affected mitochondrial morphology and not respiration (Supp. Fig. 4B-D)^32^. [U-^13^C]- palmitate tracing in 991-stimulated mDCs in presence of mdivi1 revealed that inhibition of mitochondrial fission tended to increase C_16_-labeled palmitoyl-L-carnitine abundance in 991- stimulated mDCs (Fig. 4D, E), while the abundance of palmitate-derived carbon ending up in acetyl-CoA was reduced (Fig. 4F, G). Although the contribution of palmitate to acetyl-CoA is low, the combination of increased palmitoyl-L-carnitine and lower total and labeled acetyl-CoA levels indicates lower palmitate oxidation. Together, this suggests that the AMPK-driven increase in palmitate oxidation is partly dependent on enhanced mitochondrial fission. Inhibition of mitochondrial fission did not affect tolerogenic properties of 991-stimulated mDCs (Supp. Fig. 4E, F), which may indicate that residual FAO following mdivi1 treatment is sufficient to support tolerogenic properties of the AMPK-conditioned DCs. Altogether these data indicate that AMPK activation induces mitochondrial fission to support enhanced FAO in DCs.

### Glucose oxidation contributes to AMPK-induced tolerogenicity

The increased accumulation of acetyl-CoA upon AMPK activation could only partly be explained by increased oxidation of palmitate (Fig. 3G). An important alternative carbon source for mitochondrial acetyl-CoA is glucose. To test whether increased glycolysis contributed to elevated levels of acetyl-CoA in 991-stimulated DCs, we performed [U-^13^C]-glucose tracing experiments. This revealed that AMPK activation induced a significant increase in flux of glucose-derived carbons into glucose-6- phosphate and lactate, resulting in a 3-fold and 2-fold increase in total glucose-6-phosphate and lactate levels, respectively (Fig. 5A and Supp. Fig. 5A,B), suggesting that AMPK activation enhances glycolytic flux. AMPK activation also promoted pyruvate import into the mitochondria, as increased abundance of labeled carbons in acetyl-CoA was observed (Fig. 5B and Supp. Fig. 5C,D), that could account for the increased accumulation of acetyl-CoA in these cells. The increase in glucose-derived carbon flux into acetyl-CoA in response to 991 treatment and the corresponding increase in total acetyl-CoA abundance was still partly observed in (iso)citrate, but not in further downstream TCA cycle metabolites (Fig. 5B and Supp. Fig. 5C,D). Of note, in contrast to the AMPK-induced increase in FAO, glucose oxidation was not dependent on mitochondrial fission (Supp. Fig. 4G,H). To address whether mitochondrial glucose catabolism is required for AMPK-induced tolerogenicity, we blocked mitochondrial pyruvate import using UK5099, an inhibitor for mitochondrial pyruvate carrier 1 (MPC1)^33^. While MPC1 inhibition did not restore expression of activation markers or IL-12p40 and IL- 10 secretion by 991-stimulated mDCs (Supp. Fig. 5E,F), and had mixed effects on RALDH activity, it prevented the ability of these cells to promote functional Tregs (Fig. 5C,D). Altogether, our data suggest that in addition to FAO, AMPK activation promotes mitochondrial catabolism of glucose which also underpins the tolerogenic capacity of 991-stimulated moDCs.

**Figure 5:**
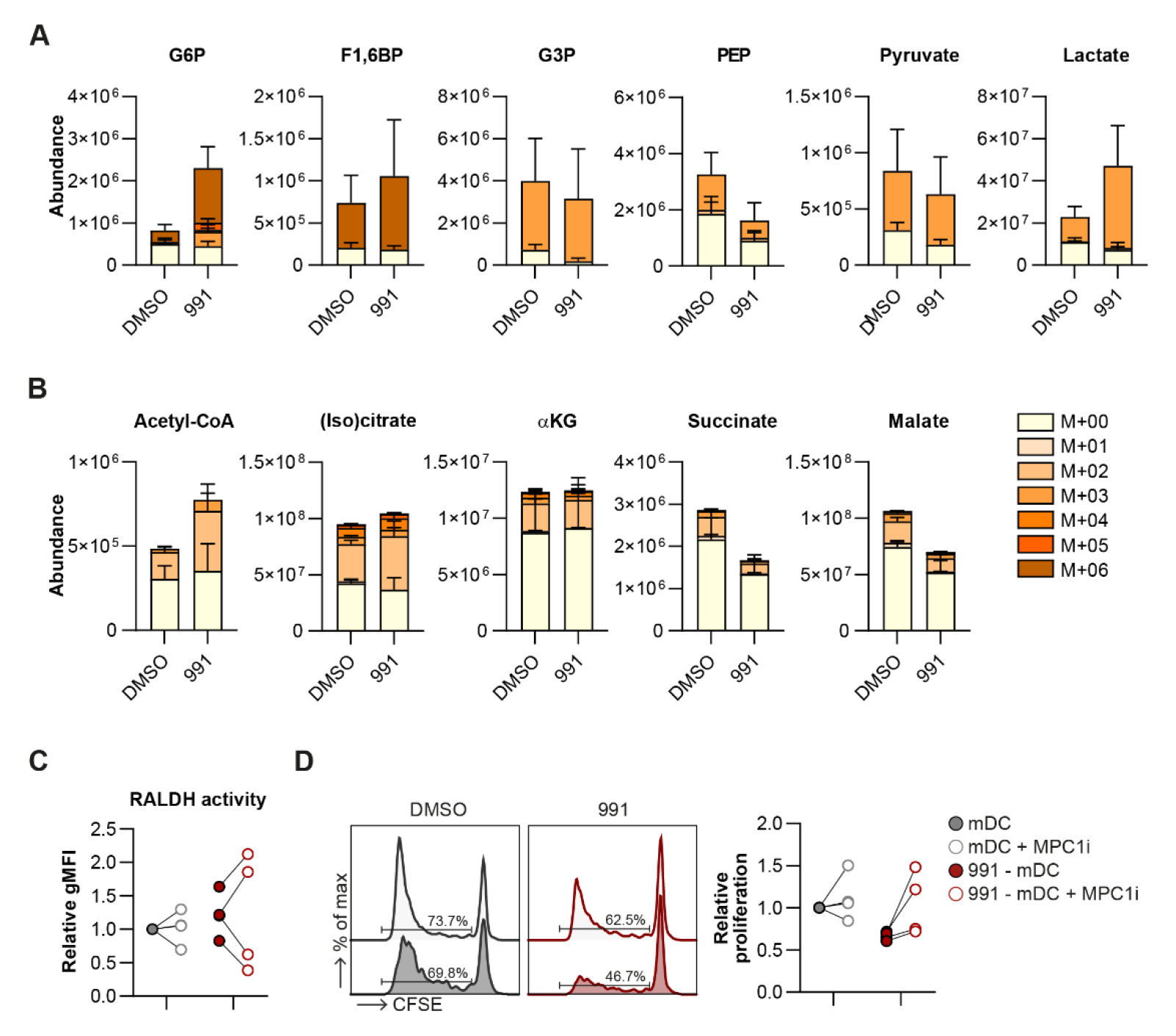
Glucose metabolism promotes AMPK-induced tolerogenicity. **A, B:** [U-^13^C]-glucose tracing was performed in DMSO/991-treated mDCs. Figures show total abundance of (labeled) metabolites involved in **(A)** glycolysis and **(B)** TCA cycle. **C:** Normalized quantification of RALDH activity. **D:** Representative histograms and normalized percentage of proliferation of bystander T cells after a T cell suppression assay. Glucose 6-phosphate (G6P), fructose 1,6-biphosphate (F1,6BP), phosphoenolpyruvate (PEP), alpha-ketoglutarate (αKG). Results are expressed as means ± SEM **(C,D)** or means ± SD **(A,B)**. Datapoints represent independent experiments with different donors. Statistical analyses were performed using two-way Anova with Tukey post-hoc test **(C, D)**. *p < 0.05, **p < 0.01.

## Discussion

Catabolic metabolism has been a well-described feature of tolDCs^9,10^. While AMPK functions as central regulator of catabolic metabolism, little is known about the role of AMPK in driving DC tolerogenicity. Despite several reports showing a role for AMPK in dampening DC activation^7,14,15^, only recently a role for AMPK in promoting regulatory T cell responses was described, as it was shown that RA-induced tolDCs require AMPK to prime Tregs^16^. On the contrary, AMPK is also activated in vitamin D3 (vitD) induced tolDCs^34^, but not required for tolerogenicity in this context^16,35^. Here, we show that direct activation of AMPK is sufficient to render DCs tolerogenic. Mechanistically, we demonstrate that this tolerogenic capacity is underpinned by enhanced RALDH activity and dependent on metabolic rewiring involving increased breakdown of glycerophospholipids, mitochondrial fission-dependent FAO, and elevated glucose catabolism.

Well-described mechanisms through which DCs induce Treg responses include IL-10^36^ or TGF- β secretion^37^, indoleamine 2,3-dioxygenase (IDO) expression^38^, and RALDH activity^39^. Our data indicate that AMPK-induced tolDCs promote Treg priming through enhanced RALDH activity. RALDH is the rate-limiting enzyme catalyzing the conversion of vitamin A-derivative retinol to RA and its expression in intestinal^19,20^ and skin DCs^40^ is important to maintain immune tolerance. We observed that inhibition of RALDH activity during moDC-T cell co-culture, but not during moDC culture alone, reduced Treg induction, suggesting that RALDH-produced RA directly acts on T cells for Treg induction.

The tolerogenic effects of AMPK activation in mDCs result in functional Tregs that do not seem to share a phenotype with any canonical Treg subset. The abundance of FoxP3^+^ CTLA4^+^ induced Tregs was lower in T cells upon priming with 991-stimulated DCs and so were ICOS^+^ Tregs, TIGIT^+^ Tregs and IL-10-secreting CD49b^+^LAG3^+^ Treg1s^21–24^. Further studies are warranted to identify the effector mechanism(s) through which Tregs primed with 991-stimulated DCs exert their immunosuppressive effects^41^.

Through an integrated proteomics and metabolomics analysis we identified breakdown of glycerophospholipids as one of the main metabolic consequence of AMPK activation in DCs. Importantly, 991-stimulated mDCs in which this process was partly inhibited through downregulation of *PNPLA6* and *GPCPD1*, were functionally less tolerogenic, demonstrating a role for glycerophospholipid breakdown in driving the tolerogenic-capacity of these moDCs. Interestingly, degradation of phospholipids has previously been reported in VitD, dexamethansone (Dex) and VitD+Dex-induced tolDCs^42^, although the functional relevance was not explored. We now show a key role for phospholipid breakdown in supporting the tolerogenic properties of DCs. Yet, how this process mechanistically underpins tolDC function remains to be addressed. Since we found the tolerogenic effects to be dependent on FAO as well, it is tempting to speculate that phospholipid catabolism supports tolDC function by means of generating free fatty acids for mitochondrial FAO. In addition, it is conceivable that lipid-mediators derived from AMPK-induced phospholipid degradation, may also exert tolerogenic effects through autocrine or paracrine mechanisms. For instance, phospholipid composition is dynamically regulated in immune cells and has been shown to govern immune cell activation through changes in membrane fluidity and production of soluble lipid mediators that mediate signal transduction and cell-cell communication^43^. Moreover, deregulation of phospholipid metabolism in tumor cells leads to accumulation of anti-inflammatory lipid mediators that suppress the anti-tumor immune response^44^. Further studies are warranted to explore whether these mechanisms also play a role in AMPK-activation driven DC tolerogenicity.

In addition to a role for glycerophospholipid catabolism, we show that AMPK activation-induced tolerogenicity is underpinned by increased FAO. We find this increase to be dependent on mitochondrial fission, a functional connection that has been reported previously in HeLa cells and which we here extend to DCs^28^. AMPK phosphorylates multiple proteins involved in mitochondrial dynamics and the observed specific increase in AMPK-mediated phosphorylation of MFF-S172/S146 suggests that AMPK activation in moDCs results in phosphorylation of fission-promoting targets, thereby inducing mitochondrial fragmentation^13^. Despite the observation that blocking FAO through CPT1A silencing compromises AMPK-induced tolerogenicity, we found that lowering FAO to baseline levels through inhibition of mitochondrial fission was not sufficient to prevent Treg induction. This indicates that the remaining levels of FAO, possibly together with elevated glucose catabolism, are sufficient to support the tolerogenic effect induced by AMPK activation. Our findings are largely consistent with earlier work showing that tolDCs induced by a combination of VitD and Dex display increased FAO, which was found to contribute to suppression of early T cell activation^8^. Moreover, tumor-associated murine DCs harbor FAO-dependent immunosuppressive capacities. Tumor-derived Wnt3a induces IDO expression and lowers IL-6 and IL-12 secretion via FAO^45^, and BMDCs treated with tumor-derived vesicles showed immune dysfunction in a PPAR-α and FAO dependent manner^46^. However, in other DC subsets FAO has also been documented to support pro-inflammatory functions, as plasmacytoid DC activation upon stimulation of endosomal toll-like-receptors was found to rely on FAO^47–49^, highlighting that the nature of the role of FAO in supporting DC function is likely DC subset dependent.

While these studies support a role for FAO in DC tolerogenicity, little is known about the mechanisms downstream of FAO that contribute to expression of anti-inflammatory genes and other immune suppressive mediators. Our data indicate that FAO mediates increased RALDH activity upon AMPK activation, although how exactly increased FAO supports RALDH activity remains unexplored. Supplementation of butyrate to moDCs is known to promote RALDH partly through inhibition of histone deacetylases (HDACs)^50^. Since AMPK can inhibit HDACs through direct phosphorylation^51^, and given that FAO-derived β-hydroxybutyrate, similar to butyrate, can inhibit HDACs^52^, it is tempting to speculate that the AMPK-induced increase in FAO together with the direct phosphorylation of the protein may contribute to HDAC inhibition and subsequent promotion of RALDH activity.

Alongside FAO, AMPK activation increased glucose catabolism as revealed by enhanced glucose-derived carbon flux into acetyl-CoA. Blocking mitochondrial pyruvate import prevented the ability of 991-stimulated moDCs to prime functional Tregs, indicating an important role for glucose-derived mitochondrial metabolites in inducing immune suppression. Increased glycolysis was also observed in DCs rendered tolerogenic by VitD alone^35,53^, VitD and Dex^34^, or by Nuclear Receptor Corepressor 1 (nCoR1) ablation^54^. While initial reports suggested that glucose availability and glycolysis, but not glucose oxidation, are essential for VitD-induced tolDCs^35^, more recent studies also indicated an essential role for glucose oxidation in VitD-induced tolerogenicity^53^. This is different from nCoR1-deficient DCs, in which MPC1 inhibition further increased tolerogenicity^54^, indicating that mitochondrial glucose oxidation may serve different roles depending on how DCs are rendered tolerogenic. In our study, we observed reduced tolerogenicity upon MPC1 inhibition, but why this is the case remains to be addressed. The relatively large contribution of glucose-derived carbons to acetyl-CoA compared to downstream TCA cycle intermediates upon AMPK activation, may point towards a role for acetyl-CoA distinct from fueling the TCA cycle in supporting AMPK-activation driven DC tolerogenicity. Acetyl-CoA can also be used for histone acetylation and in other myeloid cells, such as IL-4 polarized macrophages, acetyl-CoA accumulation fueled by glucose-derived carbons was found to be required for gene-specific histone modifications that induce M2-activation^55^. Moreover, it was recently found in T cells that glucose-derived pyruvate can be converted into acetyl-CoA in the nucleus by nuclear pyruvate dehydrogenase to alter epigenetic marks^56^. As our tracing experiments cannot differentiate between subcellular pools of acetyl-CoA, it would be worth exploring whether this is also occurring in DCs and whether the accumulation of acetyl-CoA, potentially in combination with FAO-driven effects on HDAC activity, impact epigenetic modifications in these tolerogenic DCs and thereby their function.

Overall, based on our data we propose a model in which AMPK activation promotes RALDH activity and rewiring of lipid and glucose metabolism, to render DC tolerogenic. Currently little is known about AMPK activation in DCs *in situ*, but there is some evidence that suggests this pathway is activated and may be functionally relevant in physiological tolerance promoting environments like the intestine^16^, or in pathological immunosuppressive conditions such as the tumor micro-environment^57^. Specifically, in the former setting, tolerogenic CD103^+^CD11b^+^ DCs in the murine small intestine have been shown to exhibit high AMPK activity and high RALDH activity, and AMPK deficiency in these CD11c^+^ cells induced a switch from a tolerogenic towards a more immunogenic phenotype. AMPK activation in tumor-associated DCs has, to our knowledge, not been directly quantified. However, since elevated AMPK levels have been observed in other tumor-infiltrating immune cells^58,59^ and phosphorylation of AMPK activator LKB1 has been reported in tumor-infiltrating DCs^60^, it is likely that the tumor microenvironment is also a setting with increased AMPK activation in DCs. Further studies are required to determine whether AMPK signaling or downstream pathways are relevant for DC tolerogenicity in these immunosuppressive environments. Finally, unraveling the metabolic underpinnings behind DC tolerogenicity may also provide new insights into how DC-based therapies could be improved. There have been several clinical trials in which the use of *ex vivo* or *in vivo* generated tolDCs has been evaluated for treatment of autoimmune diseases, Crohn’s disease, or to prevent transplant rejection^2–4^, unfortunately thus far, with variable success rates. Therefore, our work does not only provide new fundamental insights into the metabolic underpinnings of DC-driven tolerogenic responses, but also provides a rationale to explore AMPK as a potential actionable target for the generation of tolDCs for clinical purposes.

## Material and methods

### Monocyte-derived dendritic cell culture and stimulation

Monocytes were purified from buffy coats of healthy donors (Sanquin, Amsterdam, The Netherlands), through peripheral blood mononuclear cell isolation by Ficoll density gradient centrifugation and magnetic separation using CD14 MicroBeads (130-050-201, Miltenyi Biotec), according to manufacturer’s protocol. Monocytes were cultured in complete RPMI (RPMI-1640 (42401-042, Gibco) with 10% FCS (Greiner Bio-one), 100 U/mL penicillin (Eureco-Pharma), 100 μg/mL streptomycin (S9137, Sigma-Aldrich), and 2 mM L-glutamine (G8540, Sigma-Aldrich) supplemented with 10 ng/mL human rGM-CSF (PHC2013, Gibco) and 0.86 ng/mL human rIL4 (204-IL, R&D Systems) to induce moDC differentiation. On day 2/3 medium was refreshed and new cytokines were added. On day 5 moDCs were harvested and replated and if indicated treated with 100µM 991 (129739-36-2, Spirochem), 10 µm Mdivi1 (M0199, Sigma-Aldrich), 10 nM Vitamin D3 (D1530, Sigma-Aldrich), 45 µm Bisdiamine (WIN 18446, 4736/50, Bio-techne), 50 µm UK-5099 (PZ0160, Sigma Aldrich) or DMSO (102931, Merck Millipore). On day 6, moDCs were stimulated with 100 ng/mL ultrapure LPS (E. coli 0111 B4 strain, InvivoGen, San Diego) or left untreated. On day 7 moDCs were harvested and used for further experiments, including analysis of activation markers by flow cytometry and IL-12(p70) and IL-10 secretion by ELISA (555183 and 555157, BD Biosciences). For this, supernatant of moDCs was collected before harvesting moDCs at day 7, or after a 24 hour co-culture with CD40L-expressing J558 cells.

### T cell polarization

moDCs were cultured with allogeneic naïve CD4^+^ T cells in complete RPMI, supplemented with staphylococcal enterotoxin B (10 pg/mL). On days 6 and 8, 10 U/mL human rIL-2 (202-IL, R&D Systems) was added to expand the T cells. On day 11 cells were harvested and replated for cytokine analysis. Intracellular cytokine production was analyzed using flow cytometry after 4 hours of polyclonal restimulation with 100 ng/mL phorbol myristate acetate (PMA, P8139, Sigma-Aldrich) and 1 µg/mL ionomycin (I0634, Sigma Aldrich) in the presence of 10 µg/mL Brefeldin A (B7651, Sigma-Aldrich). IL- 10 secretion was determined by ELISA in supernatants harvested after a 24 hour restimulation with anti-CD3 (555336, BD Biosciences) and anti-CD28 (555725, BD Biosciences).

### T cell suppression assay

moDCs were cultured with allogenic naïve CD4^+^ T cells in a 1:10 ratio in complete IMDM (IMDM (BE12- 722F, Lonza) with 100 U/mL penicillin, 100 μg/mL streptomycin, 2 mM L-glutamine, and 1 mM pyruvate (P5280, Sigma-Aldrich)). On day 7 T cells (test T cells) were harvested, stained with 1uM Cell Trace Violet (C34557, Thermo Fisher Scientific), irradiated (3000 RAD), and co-cultured with autologous carboxyfluorescein succinimidyl ester (CFSE, C34554, Thermo Fisher Scientific) stained CD4^+^ memory T cells (bystander T cells) in a 2:1 ratio in the presence of LPS matured moDCs. After 6 days cells were harvested and proliferation of bystander T cells was measured using flow cytometry.

### Flow cytometry

In general, cells were stained with a viability dye (Zombie NIR™ Fixable Viability Kit, #423106 BioLegend or LIVE/DEAD™ Fixable Aqua, #L34957, Thermo) for 20 minutes at room temperature before fixation with 1.85% formaldehyde (F1635, Sigma). For assessment of phosphorylated proteins cells were fixed in 4% ultra-pure formaldehyde (#18814-20, Polysciences) for 10 minutes at 37°C directly after cell-culture, and for Treg characterization cells were fixed with FoxP3/Transcription factor staining buffer set (00-5523-00, Invitrogen) for 1 hour at 4°C. Metabolic dyes and T cell proliferation were measured in live cells. Extracellular proteins were stained for 30 minutes at 4°C in FACS buffer (PBS (220/12257974/1110, Braun), supplemented with 0.5% BSA (10735086001, Roche) and 2 mM EDTA (15575-038, Thermo Fisher Scientific). For intracellular protein and cytokine detection, cells were permeabilized for 20 minutes at 4°C in permeabilization buffer (#00-8333-56, Thermo Fisher Scientific), before staining for 30 minutes at 4°C in permeabilization buffer. Prior to staining with antibodies against phosphorylated proteins, cells were also permeabilized in 96% methanol (67-56-1, Fisher chemical) for 20 minutes at -20°C. Cells were stained with metabolic dyes as previously described^61^. Briefly, live cells were incubated with, BODIPYTM FL C16 (D3821, Thermo Fisher Scientific), BODIPYTM 493/503 (D3922, Thermo Fisher Scientific), TMRM (T668, Thermo Fisher Scientific) or MitoTracker™ Green FM (M7514, Thermo Fisher Scientific) in PBS for 15 minutes at 37°C before measurement. Aldefluor kit (01700, Stemcell Technologies) was used for assessing RALDH activity, according to manufacturer’s protocol. Briefly, cells were stained for 30 minutes at 37°C with 1 µM of Aldefluor reagent dissolved in assay buffer. Cells were kept in assay buffer until measurement. Samples were acquired on FACSCanto II or Cytek Aurora 3-laser spectral flow cytometer and analyzed using FlowJo (Version 10.6, TreeStar). Antibody information is provided in Table 1.

**Table 1:**
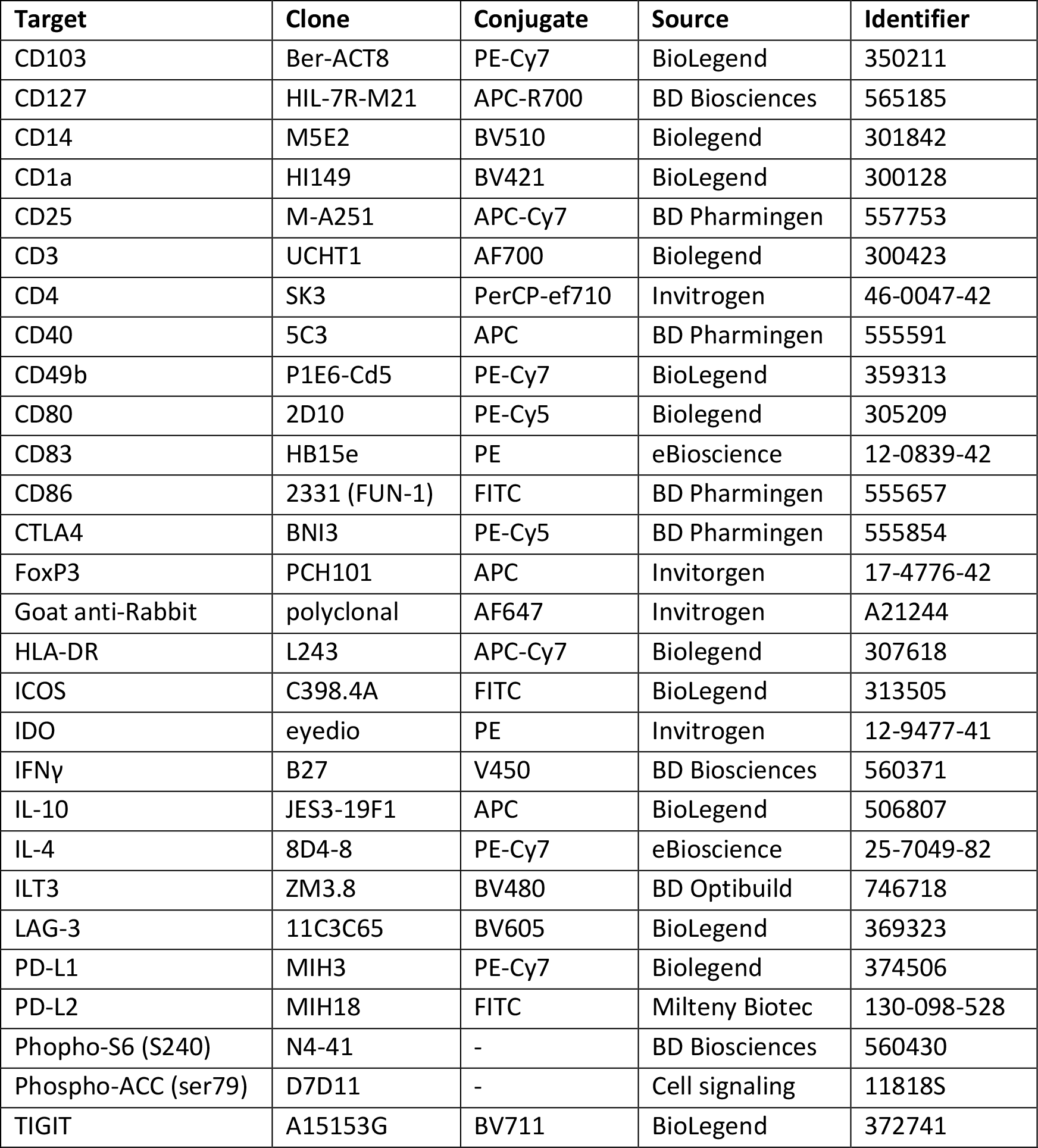
antibodies flow cytometry

### Small interfering RNA (siRNA) transfection

On day 4 of the moDC differentiation, cells were harvested and transfected with 20 nM non-targeting siRNA (D-001206-13-05, Dharmacon), or siRNAs targeting CPT1a (M-009749-02, Dharmacon), PNPLA6 (s21441, Thermo Fisher Scientific), or GPCPD1 (L-013836-01-0005, Dharmacon). Electroporation was performed with the Neon Transfection System (Invitrogen), using one pulse (1600 V, 20 ms). After electroporation, cells were overnight cultured without antibiotics and polarizing cytokines in RPMI with 10% FCS and 2 mM L-glutamine. The next morning fresh medium containing antibiotics, IL-4, and GM-CSF was added to the cells.

### Metabolomics data and analysis

Metabolites were extracted from 5×10^5^ moDCs. Cells were washed with 75 mM ammonium carbonate (A9516, Sigma-Aldrich) in HPLC-grade water (15651400, Honeywell) at 37°C, before metabolite extraction for 3 minutes in 600 µl 70% ethanol at 70°C. Samples were centrifuged at 14000 rpm for 10 min at 4°C and supernatants were shipped to General Metabolics for analysis as previously described^62^. Briefly, polar metabolites were analyzed on a non-targeted metabolomics platform in negative ion mode. 886 unique ions were identified of which 632 were annotated according to the human KEGG database. Data filtering based on interquartile range, log transformation, and normalization (mean-centered and divided by the standard deviation) was done using Metaboanalyst (www.metaboanalyst.ca), as was further analysis including pathway enrichment and differential expression analysis.

### Proteomics data and analysis

Proteins were extracted from 1×10^6^ moDCs, after cells were washed in PBS and snap-frozen in liquid nitrogen. Cell lysis, digestion and TMT labeling were performed as previously described ^63^. Cells were lysed using a 5% SDS lysis buffer (100 mM Tris-HCl pH7.6) and were incubated at 95 °C for 4 minutes. Protein determination was performed using Pierce BCA Gold protein assay (Thermo Fisher Scientific). 100 µg protein of each sample was reduced, alkylated and excess iodoacetamide quenched using with 5 mM TCEP, 15 mM iodoacetamide and 10 mM DTT, respectively. Protein was retrieved using chloroform/methanol precipitation and resulting pellets were re-solubilized in 40 mM HEPES pH 8.4. TPCK treated trypsin (1:12.5 enzyme/protein ratio) was added to digest protein overnight at 37 °C. Pierce BCA Gold protein assay was used to determine the peptide concentration. The peptides were labeled with TMTpro Label Reagents (Thermo Fisher Scientific) in a 1:4 ratio by mass (peptides/TMT reagents) for 1 h at RT. 5 µL 6% hydroxylamine was added to quench excess TMT reagent and incubated for 15 min at RT. Samples were pooled and lyophilized. The sample was subsequently fractionated on an Agilent 1200 series HPLC system (Agilent Technologies). 60 µg of the pooled peptide sample was dissolved in solvent A (10 mM NH4HCO3 pH 8.4), injected onto and eluted from an Agilent Eclipse Plus C18 2.1 x 150 mm 3.5 micron column (Agilent Technologies). The gradient was run from 2% to 90% solvent B (10 mM NH4HCO3 pH 8.4 final conc. 20/80 water/acetonitrile v/v) in 30 min at a flowrate of 200 µl/min. 12 fractions were made; every 30 sec a fraction was collected in a vial before going to the next vial. After reaching the last vial collection was continued in the first vial. Afterwards the samples were lyophilized.

Mass spectrometry: The lyophilized fractions were dissolved in water/formic acid (100/0.1 v/v) and subsequently analyzed twice by on-line C18 nanoHPLC MS/MS with a system consisting of an Easy nLC 1200 gradient HPLC system (Thermo, Bremen, Germany), and a LUMOS mass spectrometer (Thermo). Samples were injected onto a homemade precolumn (100 μm × 15 mm; Reprosil-Pur C18- AQ 3 μm, Dr. Maisch, Ammerbuch, Germany) and eluted via a homemade analytical nano-HPLC column (30 cm × 50 μm; Reprosil-Pur C18-AQ 3 um). The gradient was run from 5% to 30% solvent B (20/80/0.1 water/acetonitrile/formic acid (FA) v/v) in 240 min. The nano-HPLC column was drawn to a tip of ∼10 μm and acted as the electrospray needle of the MS source. The mass spectrometer was operated in data-dependent MS/MS mode (SPS-MS3 method) for a cycle time of 2.5 seconds, with a HCD collision energy at 36 V and recording of the MS2 spectrum in the orbitrap, with a quadrupole isolation width of 0.7 Da. In the master scan (MS1) the resolution was 120,000, the scan range 400- 1600, at standard AGC target ‘standard’. A lock mass correction on the background ion m/z=445.12 was used. Precursors were dynamically excluded after n=1 with an exclusion duration of 45 s, and with a precursor range of 20 ppm. Charge states 2-6 were included. MS2 (CID) was performed in the ion trap at 35%, with AGC target ‘standard’ and mass range ‘normal’. Synchronous precursor selection in MS3 was ‘true’ at a HCD collision energy of 55% and detection in the orbitrap at a resolution of 50,000, at an AGC target of 300% with a scan range of m/z 100-500. In a post-analysis process, raw data were first converted to peak lists using Proteome Discoverer version 2.5 (Thermo Electron), and submitted to the Uniprot database (Homo sapiens, 20596 entries), using Mascot v. 2.2.07 (www.matrixscience.com) for protein identification. Mascot searches were with 10 ppm and 0.5 Da deviation for precursor and fragment mass, respectively. Enzyme specificity was set to trypsin. Up to three missed cleavages were allowed. Methionine oxidation and acetyl on protein N-terminus were set as variable modifications. Carbamidomethyl on Cys and TMTpro on Lys and N-terminus were set as fixed modifications. Protein FDR of 1% was set. Normalization was on total peptide amount. 8325 proteins were identified of which 3462 were used for further processing after filtering based on high protein confidence and at least 5 unique peptides per protein. Pathway enrichment was performed using Metascape and enriched gene ontology (GO) pathways were considered significant if FDR ≤ 0.05 ^64^. Integration of metabolomics and proteomics data was performed using Shiny Gam (https://artyomovlab.wustl.edu/shiny/gam/)^25^.

### ^13^C-tracer experiments

On day 7, 20 hours after LPS stimulation, moDCs were harvested, counted and 5×10^5^ cells were replated in 350 µl in complete glucose-free RPMI (11879020, Thermo Fisher Scientific), supplemented with 100 U/mL penicillin, 100 μg/mL streptomycin, and 25 mM HEPES (9136, Sigma Aldrich). Additionally, 350 µl complete glucose-free RPMI was added, supplemented with either 22 mM [U-^13^C]- Glucose (CLM-1396-PK, Cambridge Isotope Laboratories) and 0.027 mM ^12^C-Palmitate (76119, Merck Millipore) or 22 mM ^12^C-Glucose (15023021, Gibco) and 0.027 mM [U-^13^C]-Palmitate (CLM-409-PK, Cambridge Isotope Laboraties), to reach an end concentration of 11 mM glucose and 0.014 mM palmitate. Palmitate was in-house conjugated to BSA in a 2:1 ratio. Cells were cultured for 4 hours at 37°C before metabolites were extracted. Cells were washed in ice-cold PBS, before extraction in lysis buffer (40% acetonitrile (34967, Honeywell) and 40% methanol in HPLC-grade water). Samples were vortexed 3 times for 3 seconds and centrifuged at 4°C for 15 minutes at full speed. Supernatant was stored at -80°C. Metabolites were analyzed by liquid chromatography – mass spectrometry (LC-MS)- based metabolomics as previously described^65^. Briefly, metabolite identification was based on exact mass (5 ppm range), validated by concordance with retention times of standards and peak areas were in their respective linear range of detection. TraceFinder Software (Thermo Fisher Scientific) was used for quantification. Peak intensities were normalized based on mean peak intensity and isotopomer distributions were corrected for natural abundance.

### Quantitative polymerase chain reaction (qPCR)

RNA was extracted from snap-frozen moDCs as previously described (^16^). cDNA synthesis was performed using M-MLV Reverse Transcriptase (28025013, Thermo Fisher Scientific) and qPCR was performed with the Gotaq qPCR master mix (A6001, Promega) and the CFX96 Touch Real-Time PCR Detection System (Biorad). Primers are listed in table 2.

**Table 2:**
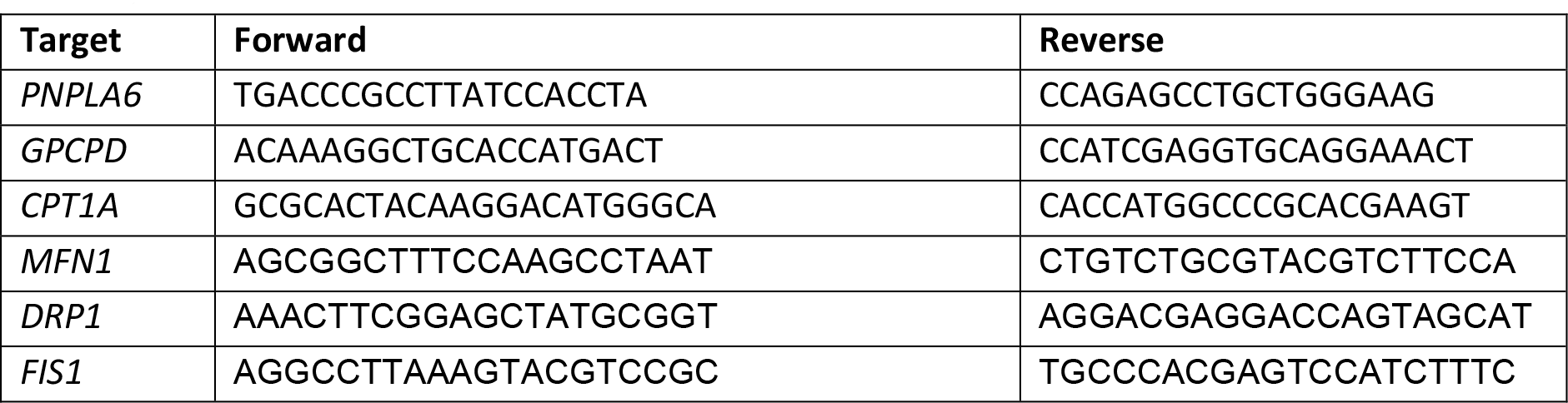
primers

### Confocal microscopy

On day 5 of moDC differentiation, cells were plated in glass bottom confocal dishes (627871, Greiner) and coated for 2 hours with Poly-D-Lysine (PDL) at 37°C. Treatments were performed as described earlier and cells were analyzed on day 7. 10 minutes prior to imaging nuclei were stained with 50 ng/mL Hoechst 33342 (H3570, Thermo Fisher Scientific) and mitochondria with 10 nM Cy5-methyl-methyl (Cy5M_2_)^66^. Cells were imaged directly using the 40x objective on a SP8X WLL (white light laser) microscope (Leica Microsystems, Wetzlar, Germany) with climate control at 37°C, 5% CO_2_. Mitochondrial quantification was done in ImageJ using the MitoAnalyzer Marco as previously described^67^.

### Seahorse extracellular flux analysis

Mitochondria oxygen consumption rate (OCR) was analyzed using a XF^e^96 extracellular flux analyzer (Seahorse Bioscience). moDCs were plated in seahorse cell culture plates coated with Poly-D-Lysine (PDL) at 37°C for 1 hour, in unbuffered, glucose-free RPMI, supplemented with 5% FCS and 2 mM L-glutamine, and incubated for 1 hour at 37°C in a CO_2_ free incubator. OCR was measured in response to 10 mM glucose (G8644, Sigma-Aldrich), 1 µM oligomycin (11342, Cayman Chemical), 3 µM fluoro-carbonyl cyanide phenylhydrazone (FCCP) (C2920-Sigma-Aldrich) and 1 µM rotenone (R8875, Sigma-Aldrich) + 1 µM antimycin A (A8674, Sigma-Aldrich).

### Western blot

moDCs were washed in PBS, snap-frozen in liquid nitrogen and lysed with EBSB buffer (8% glycerol (7044, J. T. Baker), 3% SDS (151-21-3, VWR Life Science) and 100 mM Tris–HCl [pH 6.8](1.00317, Supelco; 77-86-1, Thermo Fisher Scientific)) for detection of complexes of the electron transport chain, or lysed with RIPA-like buffer (50 mmol/L HEPES (pH 7.6), 50 mmol/L Natrium fluoride (S7920, Sigma Aldrich), 50 mmol/L Potassium chloride (7447407, Merck Millipore), 5 mmol/L Sodium pyrophosphate decahydrate (13472-36-1, Sigma Aldrich), 1 mmol/L EDTA (E5134, Sigma Aldrich), 1 mmol/L EGTA (E3889, Sigma Aldrich), 1 mmol/L dithiothreitol (V315A, Promega), 5 mmol/L β-glycerophosphate (G6376, Sigma Aldrich), 1 mmol/L sodium orthovanadate (S6508, Sigma Aldrich), 1% NP40 (74385, Sigma Aldrich), and protease inhibitor cocktail (11697498001; Roche). Lysates were boiled for 5 minutes (except lysates used for detection of electron transport chain complexes) and protein content was determined using a BCA kit (23225, Thermo Fisher Scientific). Proteins were separated by SDS-PAGE and transferred to a 45 µm PVDF membrane. Membranes were blocked for 1 hour at room temperature in TTBS buffer (20 mM Tris-HCl [pH 7.6], 137 mM NaCl (1.06404.1000, Merck Millipore), and 0.25% Tween (8.22184.0500, Merck Millipore)) containing 5% fat free milk (Campina). Membranes were incubated with primary antibodies overnight at 4°C, washed in TTBS buffer, incubated with horseradish peroxidase-conjugated secondary antibodies for 2 hours at room temperature, washed again and developed using enhanced chemiluminescence. Primary antibodies: HSP90 (Santa Cruz #sc7947), DRP1 (5391T, Cell Signaling), Phospho-DRP1 (Ser637) (4867S, Cell Signaling), MFF (84580T, Cell Signaling), Phospho-MFF (Ser172, Ser146) (PA5-104614, Thermo Fisher Scientific), total Oxphos antibody cocktail (ab110413, Abcam).

Western blots were quantified using ImageJ software (NIH, Bethesda, MD, USA) and phosphorylation status was normalized to the total abundance of the target protein and a housekeeping protein.

### Statistical analysis

Results are expressed as mean ± standard error mean (SEM) except where stated otherwise. Data were analyzed using GraphPad Prism (La Jolla, CA, USA). Comparisons between two or more independent data groups were made by Student’s T test or analysis of variance test (ANOVA),respectively. P<0.05 was considered statistically significant.

## Acknowledgements

This work was supported by an LUMC fellowship awarded to BE.

## Author Contributions

ECB and BE designed and analyzed the experiments. ECB, TAP, AJH, TJAM, and FO performed the experiments. FWMV, EAZ, and CRB aided with the ^13^C-tracer experiments. AHR, RTNT, and PAV aided with the proteomics experiments. BG helped in the discussion and reviewed the manuscript. BE conceived and supervised the study and wrote the manuscript together with ECB.

## References

1. Kapsenberg, M. L. Dendritic-cell control of pathogen-driven T-cell polarization. Nat Rev Immunol 3, 984–993 (2003).

2. Mohammadi, B. et al. The role of tolerogenic dendritic cells in systematic lupus erythematosus progression and remission. Int Immunopharmacol 115, (2023).

3. Ness, S., Lin, S. & Gordon, J. R. Regulatory Dendritic Cells, T Cell Tolerance, and Dendritic Cell Therapy for Immunologic Disease. Front Immunol 12, (2021).

4. Passeri, L., Marta, F., Bassi, V. & Gregori, S. Tolerogenic dendritic cell-based approaches in autoimmunity. Int J Mol Sci 22, (2021).

5. Brombacher, E. C. & Everts, B. Shaping of Dendritic Cell Function by the Metabolic Micro-Environment. Front Endocrinol (Lausanne) 11, (2020).

6. Everts, B. et al. TLR-driven early glycolytic reprogramming via the kinases TBK1-IKKε supports the anabolic demands of dendritic cell activation. Nat Immunol 15, 323–332 (2014).

7. Krawczyk, C. M. et al. Toll-like receptor-induced changes in glycolytic metabolism regulate dendritic cell activation. Blood 115, 4742–4749 (2010).

8. Malinarich, F. et al. High Mitochondrial Respiration and Glycolytic Capacity Represent a Metabolic Phenotype of Human Tolerogenic Dendritic Cells. The Journal of Immunology 194, 5174–5186 (2015).

9. Patente, T. A., Pelgrom, L. R. & Everts, B. Dendritic cells are what they eat: how their metabolism shapes T helper cell polarization. Curr Opin Immunol 58, 16–23 (2019).

10. Møller, S. H., Wang, L. & Ho, P. C. Metabolic programming in dendritic cells tailors immune responses and homeostasis. Cell Mol Immunol 19, 370–383 (2022).

11. Steinberg, G. R. & Hardie, D. G. New insights into activation and function of the AMPK. Nat Rev Mol Cell Biol 24, 255–272 (2022).

12. Snyder, J. P. & Amiel, E. Regulation of dendritic cell immune function and metabolism by cellular nutrient sensor mammalian target of rapamycin (mTOR). Front Immunol 10, (2019).

13. Trefts, E. & Shaw, R. J. AMPK: restoring metabolic homeostasis over space and time. Mol Cell 81, 3677–3690 (2021).

14. Carroll, K. C., Viollet, B. & Suttles, J. AMPKα1 deficiency amplifies proinflammatory myeloid APC activity and CD40 signaling. J Leukoc Biol 94, 1113–1121 (2013).

15. Nieves, W. et al. Myeloid-Restricted AMPKα1 Promotes Host Immunity and Protects against IL-12/23p40–Dependent Lung Injury during Hookworm Infection. The Journal of Immunology 196, 4632–4640 (2016).

16. Patente, T. A. et al. Metabolic sensor AMPK licenses CD103+ dendritic cells to induce Treg responses. bioRxiv (2023).

17. Bultot, L. et al. Benzimidazole derivative small-molecule 991 enhances AMPK activityand glucose uptake induced by AICAR or contraction in skeletal muscle. Am J Physiol Endocrinol Metab 311, 706–719 (2016).

18. Hardie, D. G., Ross, F. A. & Hawley, S. A. AMPK: A nutrient and energy sensor that maintains energy homeostasis. Nat Rev Mol Cell Biol 13, 251–262 (2012).

19. Sun, C. M. et al. Small intestine lamina propria dendritic cells promote de novo generation of Foxp3 T reg cells via retinoic acid. Journal of Experimental Medicine 204, 1775–1785 (2007).

20. Coombes, J. L. et al. A functionally specialized population of mucosal CD103+ DCs induces Foxp3+ regulatory T cells via a TGF-β -and retinoic acid-dependent mechanism. Journal of Experimental Medicine 204, 1757–1764 (2007).

21. Boonpiyathad, T., Sözener, Z. C., Akdis, M. & Akdis, C. A. The role of treg cell subsets in allergic disease. Asian Pac J Allergy Immunol 38, 139–149 (2020).

22. Zheng, J. et al. ICOS regulates the generation and function of human CD4+ Treg in a CTLA-4 dependent manner. PLoS One 8, (2013).

23. Joller, N. et al. Treg cells expressing the coinhibitory molecule TIGIT selectively inhibit proinflammatory Th1 and Th17 cell responses. Immunity 40, 569–581 (2014).

24. Gagliani, N. et al. Coexpression of CD49b and LAG-3 identifies human and mouse T regulatory type 1 cells. Nat Med 19, 739–746 (2013).

25. Sergushichev, A. A. et al. GAM: a web-service for integrated transcriptional and metabolic network analysis. Nucleic Acids Res 44, 194–200 (2016).

26. Van Tienhoven, M., Atkins, J., Li, Y. & Glynn, P. Human neuropathy target esterase catalyzes hydrolysis of membrane lipids. Journal of Biological Chemistry 277, 20942–20948 (2002).

27. Okazaki, Y. et al. A novel glycerophosphodiester phosphodiesterase, GDE5, controls skeletal muscle development via a non-enzymatic mechanism. Journal of Biological Chemistry 285, 27652–27663 (2010).

28. Song, J. E. et al. Mitochondrial fission governed by drp1 regulates exogenous fatty acid usage and storage in hela cells. Metabolites 11, (2021).

29. Li, J. et al. Pharmacological activation of AMPK prevents Drp1-mediated mitochondrial fission and alleviates endoplasmic reticulum stress-associated endothelial dysfunction. J Mol Cell Cardiol 86, 62–74 (2015).

30. Toyama, E. Q. et al. Metabolism: AMP-activated protein kinase mediates mitochondrial fission in response to energy stress. Science (1979) 351, 275–281 (2016).

31. Cassidy-Stone, A. et al. Chemical Inhibition of the Mitochondrial Division Dynamin Reveals Its Role in Bax/Bak-Dependent Mitochondrial Outer Membrane Permeabilization. Dev Cell 14, 193–204 (2008).

32. Bordt, E. A. et al. The Putative Drp1 Inhibitor mdivi-1 Is a Reversible Mitochondrial Complex I Inhibitor that Modulates Reactive Oxygen Species. Dev Cell 40, 583–594 (2017).

33. Halestrap, A. P. The Mitochondrial Pyruvate Carrier. Kinetics and specificity for substrated and inhibitors. Biochem. J vol. 148 (1975).

34. Adamik, J. et al. Distinct metabolic states guide maturation of inflammatory and tolerogenic dendritic cells. Nat Commun 13, 1–19 (2022).

35. Ferreira, G. B. et al. Vitamin D3 induces tolerance in human dendritic cells by activation of intracellular metabolic pathways. Cell Rep 10, 711–725 (2015).

36. Boks, M. A. et al. IL-10-generated tolerogenic dendritic cells are optimal for functional regulatory T cell induction - A comparative study of human clinical-applicable DC. Clinical Immunology 142, 332–342 (2012).

37. Awasthi, A. et al. A dominant function for interleukin 27 in generating interleukin 10- producing anti-inflammatory T cells. Nat Immunol 8, 1380–1389 (2007).

38. Rodrigues, C. P. et al. Tolerogenic IDO+ dendritic cells are induced by pd-1-expressing mast cells. Front Immunol 7, (2016).

39. Mucida Daniel et al. Reciprocal TH17 and RegulatoryT Cell Differentiation Mediated by Retinoic Acid. Science (1979) 317, 256–260 (2007).

40. Weckel, A. et al. Long-term tolerance to skin commensals is established neonatally through a specialized dendritic cell subgroup. Immunity 56, (2023).

41. Chung, D. J. et al. Indoleamine 2,3-dioxygenase-expressing mature human monocyte-derived dendritic cells expand potent autologous regulatory T cells. Blood 114, 555–563 (2009).

42. Ferreira, G. B. et al. Differential protein pathways in 1,25-dihydroxyvitamin D 3 and dexamethasone modulated tolerogenic human dendritic cells. J Proteome Res 11, 941–971 (2012).

43. O’Donnell, V. B., Rossjohn, J. & Wakelam, M. J. O. Phospholipid signaling in innate immune cells. Journal of Clinical Investigation 128, 2670–2679 (2018).

44. Saito, R. de F., Andrade, L. N. de S., Bustos, S. O. & Chammas, R. Phosphatidylcholine-Derived Lipid Mediators: The Crosstalk Between Cancer Cells and Immune Cells. Front Immunol 13, (2022).

45. Zhao, F. et al. Paracrine Wnt5a-β-Catenin Signaling Triggers a Metabolic Program that Drives Dendritic Cell Tolerization. Immunity 48, 147–160 (2018).

46. Yin, X. et al. PPARα Inhibition Overcomes Tumor-Derived Exosomal Lipid-Induced Dendritic Cell Dysfunction. Cell Rep 33, (2020).

47. Qiu, C. C., Atencio, A. E. & Gallucci, S. Inhibition of fatty acid metabolism by etomoxir or TOFA suppresses murine dendritic cell activation without affecting viability. Immunopharmacol Immunotoxicol 41, 361–369 (2019).

48. Wu, D. et al. Type 1 Interferons Induce Changes in Core Metabolism that Are Critical for Immune Function. Immunity 44, 1325–1336 (2016).

49. Basit, F. & de Vries, I. J. M. Dendritic Cells Require PINK1-Mediated Phosphorylation of BCKDE1α to Promote Fatty Acid Oxidation for Immune Function. Front Immunol 10, (2019).

50. Kaisar, M. M. M., Pelgrom, L. R., van der Ham, A. J., Yazdanbakhsh, M. & Everts, B. Butyrate conditions human dendritic cells to prime type 1 regulatory T cells via both histone deacetylase inhibition and G protein-coupled receptor 109A signaling. Front Immunol 8, (2017).

51. Mihaylova, M. M. et al. Class IIa histone deacetylases are hormone-activated regulators of FOXO and mammalian glucose homeostasis. Cell 145, 607–621 (2011).

52. Shimazu, T. et al. Suppression of oxidative stress by β-hydroxybutyrate, an endogenous histone deacetylase inhibitor. Science (1979) 339, 211–214 (2013).

53. Vanherwegen, A. S. et al. Vitamin D controls the capacity of human dendritic cells to induce functional regulatory T cells by regulation of glucose metabolism. Journal of Steroid Biochemistry and Molecular Biology 187, 134–145 (2019).

54. Sen, K. et al. NCoR1 controls immune tolerance in conventional dendritic cells by fine-tuning glycolysis and fatty acid oxidation. Redox Biol 59, (2023).

55. Covarrubias, A. J. et al. Akt-mTORC1 signaling regulates Acly to integrate metabolic input to control of macrophage activation. 5, (2016).

56. Mocholi, E. et al. Pyruvate metabolism controls chromatin remodeling during CD4+ T cell activation. Cell Rep 42, (2023).

57. Arner, E. N. & Rathmell, J. C. Metabolic programming and immune suppression in the tumor microenvironment. Cancer Cell 41, 421–433 (2023).

58. Trillo-Tinoco, J. et al. AMPK alpha-1 intrinsically regulates the function and differentiation of tumor myeloid-derived suppressor cells. Cancer Res 79, 5034–5047 (2019).

59. An, J., et al. AMP-activated protein kinase alpha1 promotes tumor development via FOXP3 elevation in tumor-infiltrating Treg cells. iScience 25, (2022).

60. Wang, Y. et al. LKB1 orchestrates dendritic cell metabolic quiescence and anti-tumor immunity. Cell Res 29, 391–405 (2019).

61. Brombacher, E. C., Patente, T. A., Quik, M. & Everts, B. Characterization of Dendritic Cell Metabolism by Flow Cytometry. in Dendritic Cells. Methods in Molecular Biology (ed. Sisirak, V.) vol. 2618 219–237 (Humana Press, 2023).

62. Fuhrer, T., Heer, D., Begemann, B. & Zamboni, N. High-throughput, accurate mass metabolome profiling of cellular extracts by flow injection-time-of-flight mass spectrometry. Anal Chem 83, 7074–7080 (2011).

63. Paulo, J. A. & Gygi, S. P. Nicotine-induced protein expression profiling reveals mutually altered proteins across four human cell lines. Proteomics 17, (2017).

64. Zhou, Y. et al. Metascape provides a biologist-oriented resource for the analysis of systems-level datasets. Nat Commun 10, (2019).

65. Zaal, E. A. et al. Bortezomib resistance in multiple myeloma is associated with increased serine synthesis. Cancer Metab 5, (2017).

66. Winkel, B. M. F. et al. A tracer-based method enables tracking of plasmodium falciparum malaria parasites during human skin infection. Theranostics 9, 2768–2778 (2019).

67. Chaudhry, A., Shi, R. & Luciani, D. S. A pipeline for multidimensional confocal analysis of mitochondrial morphology, function, and dynamics in pancreatic b-cells. Am J Physiol Endocrinol Metab 318, 87–101 (2020).

