## Supplementary figures for "AMPK activation induces RALDH^high^ tolerogenic dendritic cells through rewiring of glucose and lipid metabolism"

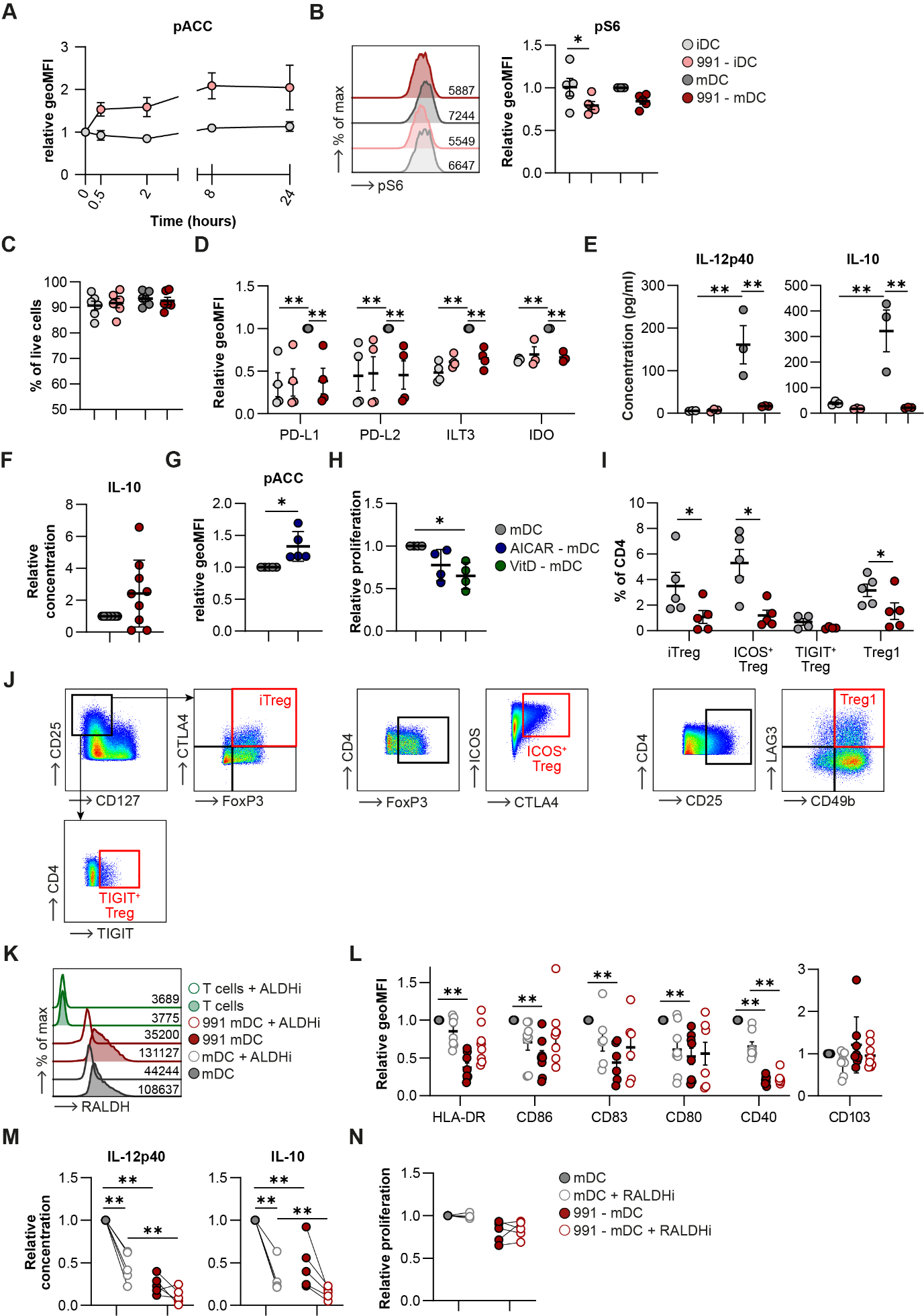

**Supplementary figure 1. Effects of 991 and RALDH inhibition on moDCs and T cell priming.**

**A:** Phosphorylation of ACC (Ser79) at different time points as determined by flow cytometry. **B:** Representative histogram and normalized quantification of phosphorylation of S6 (Ser240). **C:** Percentage of live cells as % of total moDCs. **D:** Normalized expression of indicated markers on moDCs. **E:** Concentration of IL-12p40 and IL-10 in the supernatants of moDCs. **F:** Relative concentration of IL-10 in the supernatants of T cells primed with 991/DMSO-treated mDCs after a 24 hour restimulation with αCD3/αCD28 antibodies. **G:** Normalized quantification of phosphorylation of ACC (Ser79) after treatment with H^2^O/AICAR. **H:** Normalized percentage of proliferation of bystander T cells after a T cell suppression assay. **I:** Abundance of CD25^+^FoxP3^+^CTLA4^+^ induced Tregs (iTregs), CD25^+^FoxP3^+^ICOS^+^ Tregs (ICOS^+^ Treg), CD25^+^TIGIT^+^ Treg (TIGIT^+^ Treg) and CD25^+^CD49b^+^LAG3^+^ Tregs (Treg1) in total CD4^+^ T cells after priming with DMSO/991-treated mDCs. **J:** Gating strategy of T cell subsets as described in I. **K:** representative histograms of RALDH activity after treatment with RALDH inhibitor bisdiamine (RALDHi). Representative of 4 independent experiments. **L:** Normalized expression of indicated markers. **M:** Normalized concentration of IL-12p40 and IL-10 after a 24hour co-culture with with CD40L-expressing J558 cells. **N:** Normalized quantification of bystander T cell proliferation after co-culture with irradiated T cells primed with 991/DMSO-treated mDC cultured in presence or absence of RALDH inhibitor bisdiamine (RALDHi). Results are expressed as means ± SEM. Datapoints represent independent experiments with different donors. Statistical analyses were performed using paired t-tests **(F, G, I)**, one-way Anova with Sidak post-hoc test **(H)** or two-way Anova with Tukey post-hoc test **(A-E, L-N)**. *p < 0.05, **p < 0.01.

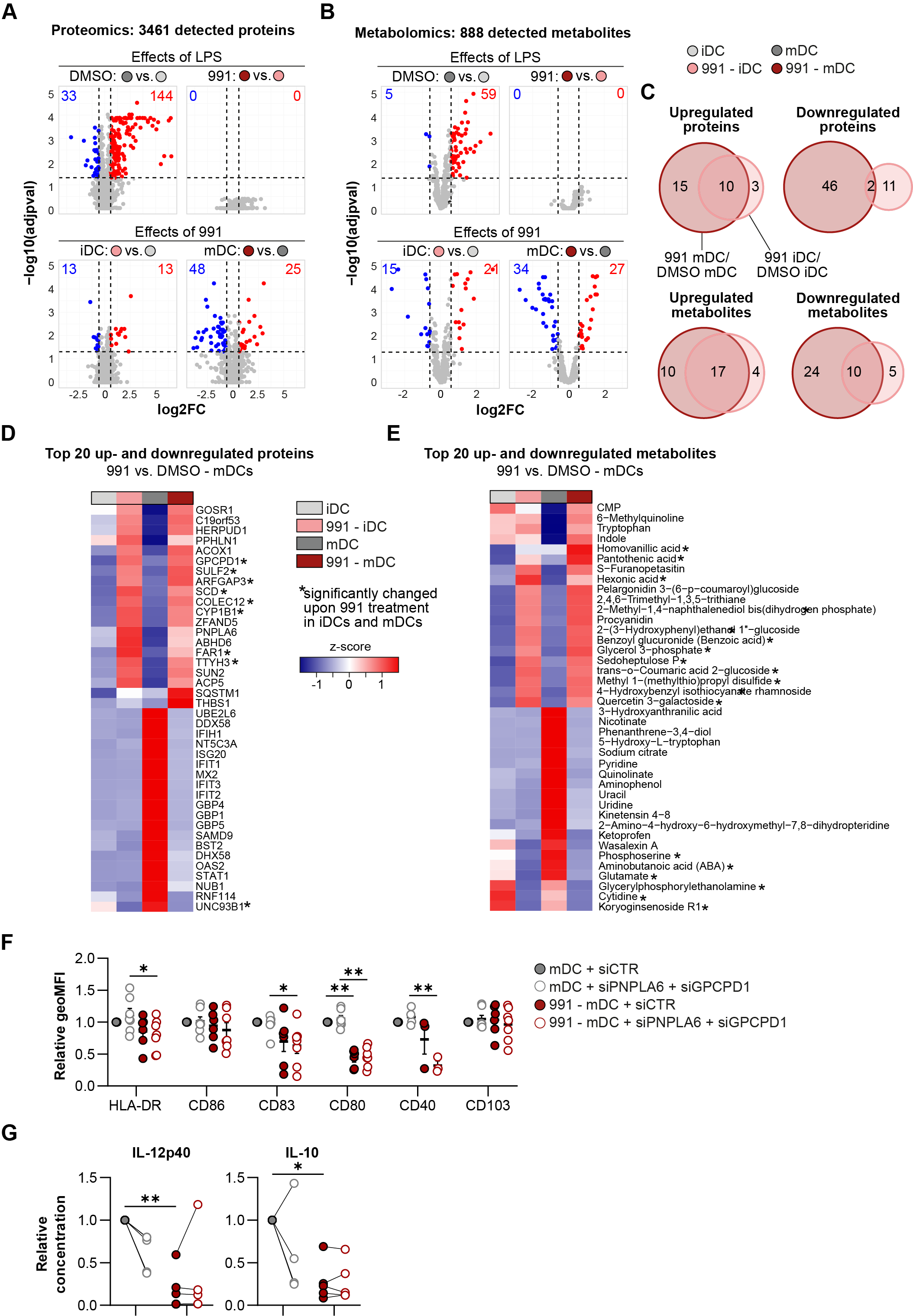

**Supplementary figure 2: Effects of AMPK activation on proteome and metabolome of moDCs.**

**A,B:** Volcano plots depicting all identified **(A)** proteins and **(B)** metabolites. Blue and red dots denote the significantly down- and upregulated proteins/metabolites, respectively. **C:** Venn diagrams showing the overlap between up- and down-regulated proteins and metabolites upon AMPK activation in mDCs and iDCs. **D, E:** Heatmaps depicting the relative expression of the 20 most up- and downregulated **(D)** proteins and **(E)** metabolites upon AMPK activation in mDCs, ranked by log2 fold change and filtered by adjusted p-value <0.05. **F:** Normalized expression of indicated markers. **G:** relative concentration of IL-12p40 and IL-10 in the supernatants of mDCs after co-culture with CD40L-expressing J558 cells. Results are expressed as means ± SEM. Datapoints represent independent experiments with different donors. Statistical analyses were performed two-way Anova with Tukey post-hoc test **(F, G)**. *p < 0.05, **p < 0.01.

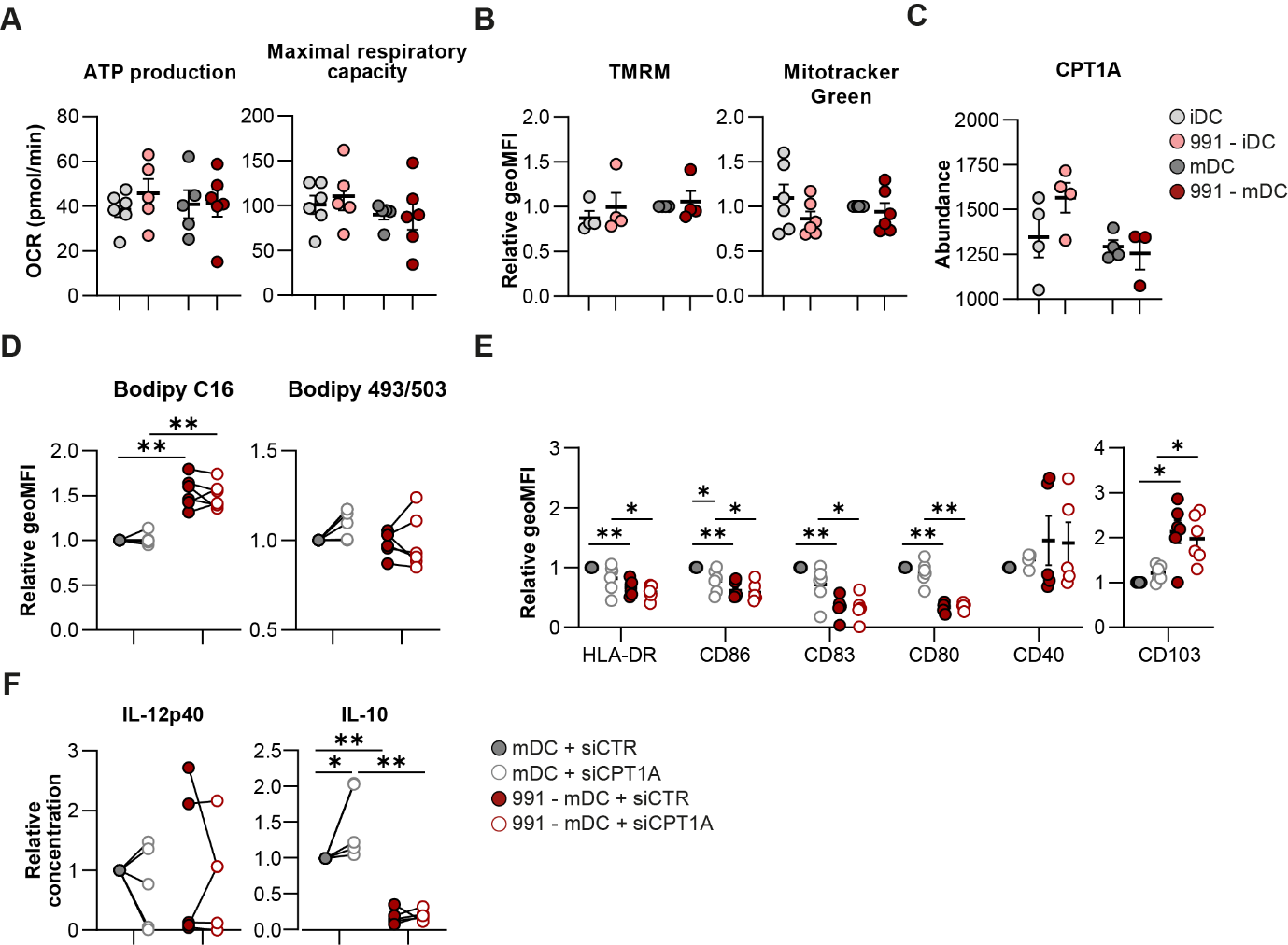

**Supplementary figure 3: effects of AMPK activation on mitochondrial metabolism.**

**A:** Quantification of ATP production and maximal respiratory capacity of oxygen consumption rate (OCR) derived from Figure 3A. **B:** Normalized quantification of mitochondrial membrane potential (TMRM) and mitochondrial mass (mitotracker green). **C:** Expression of CPT1A derived from the proteomics data. **D:** Normalized quantification of Bodipy C16 and Bodipy 493/503 staining. **E:** Normalized expression of indicated markers. **F:** Normalized concentration of IL-12p40 and IL-10 after a 24hour co-culture with with CD40L-expressing J558 cells. Results are expressed as means ± SEM. Datapoints represent independent experiments with different donors. Statistical analyses were performed using two-way Anova with Tukey post-hoc test **(A-F)**. *p < 0.05, **p < 0.01.

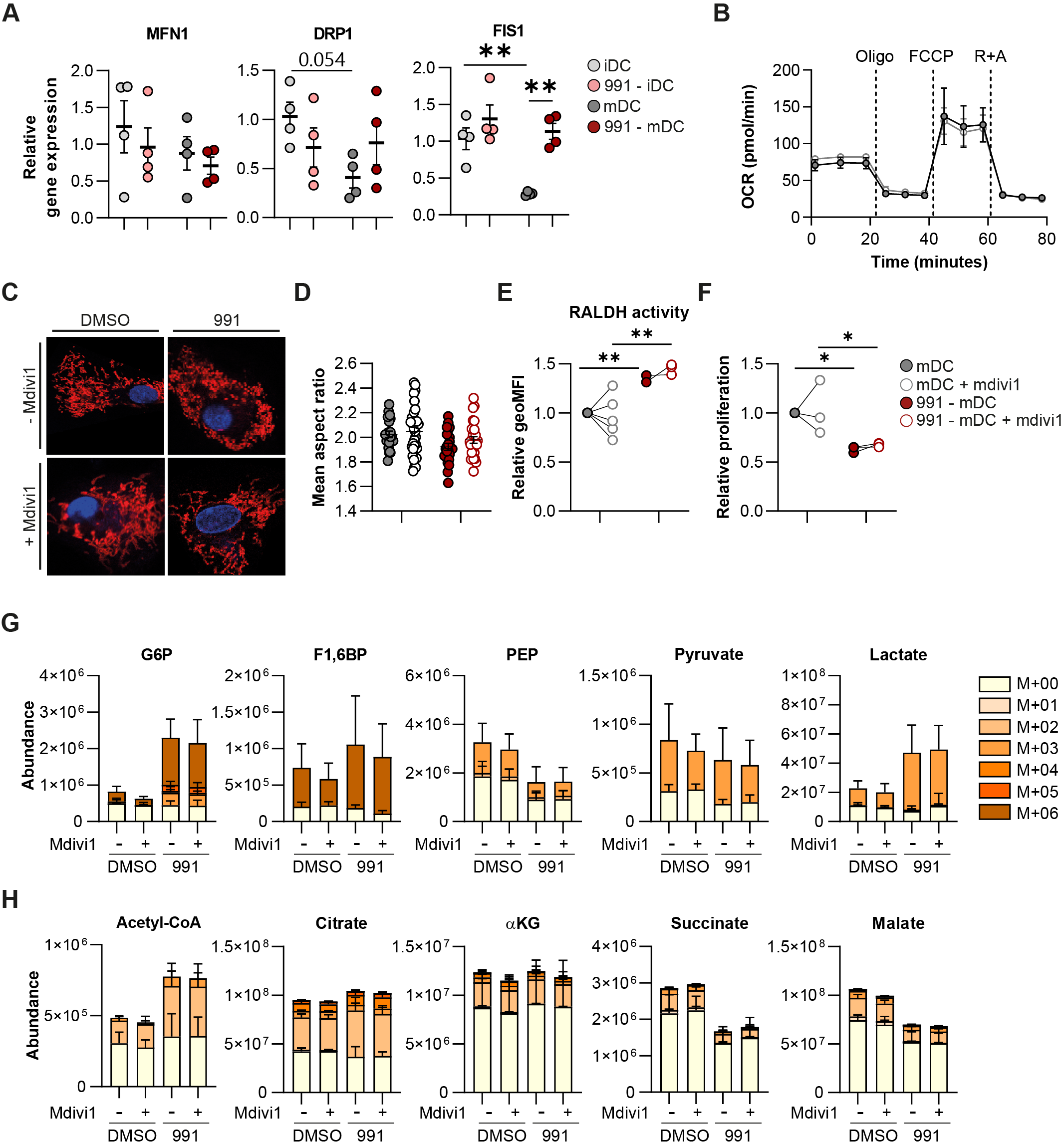

**Supplementary figure 4: effects of AMPK activation on mitochondrial fission.**

**A:** Relative gene expression of genes regulating mitochondrial dynamics. **B:** Real-time oxygen consumption rate (OCR) as measured by Seahorse extracellular flux analysis. **C:** Confocal images of DMSO/991-treated mDCs with or without mdivi1 treatment. Blue: nucleus. Red: mitochondria. **D:** Quantification of the average aspect ratio of mitochondria per cell. **E: N**ormalized quantification of RALDH activity and **(F)** normalized percentage of proliferation of bystander T cells after a T cell suppression assay. **G, H:** [U-^13^C]-glucose tracing was performed in DMSO/991-treated mDCs, with or without mdivi1. Figures show total abundance of (labeled) metabolites involved in **(G)** glycolysis and **(H)** TCA cycle. Glucose 6-phosphate (G6P), fructose 1,6-biphosphate (F1,6BP), glycerol 3-phosphate (G3P), phosphoenolpyruvate (PEP), alpha-ketoglutarate (αKG). Results are expressed as means ± SEM **(A, D,E)** or means ± SD **(F,G)**. Datapoints represent independent experiments with different donors. Statistical analyses were performed using two-way Anova with Tukey post-hoc test **(A, D-F)**. *p < 0.05, **p < 0.01.

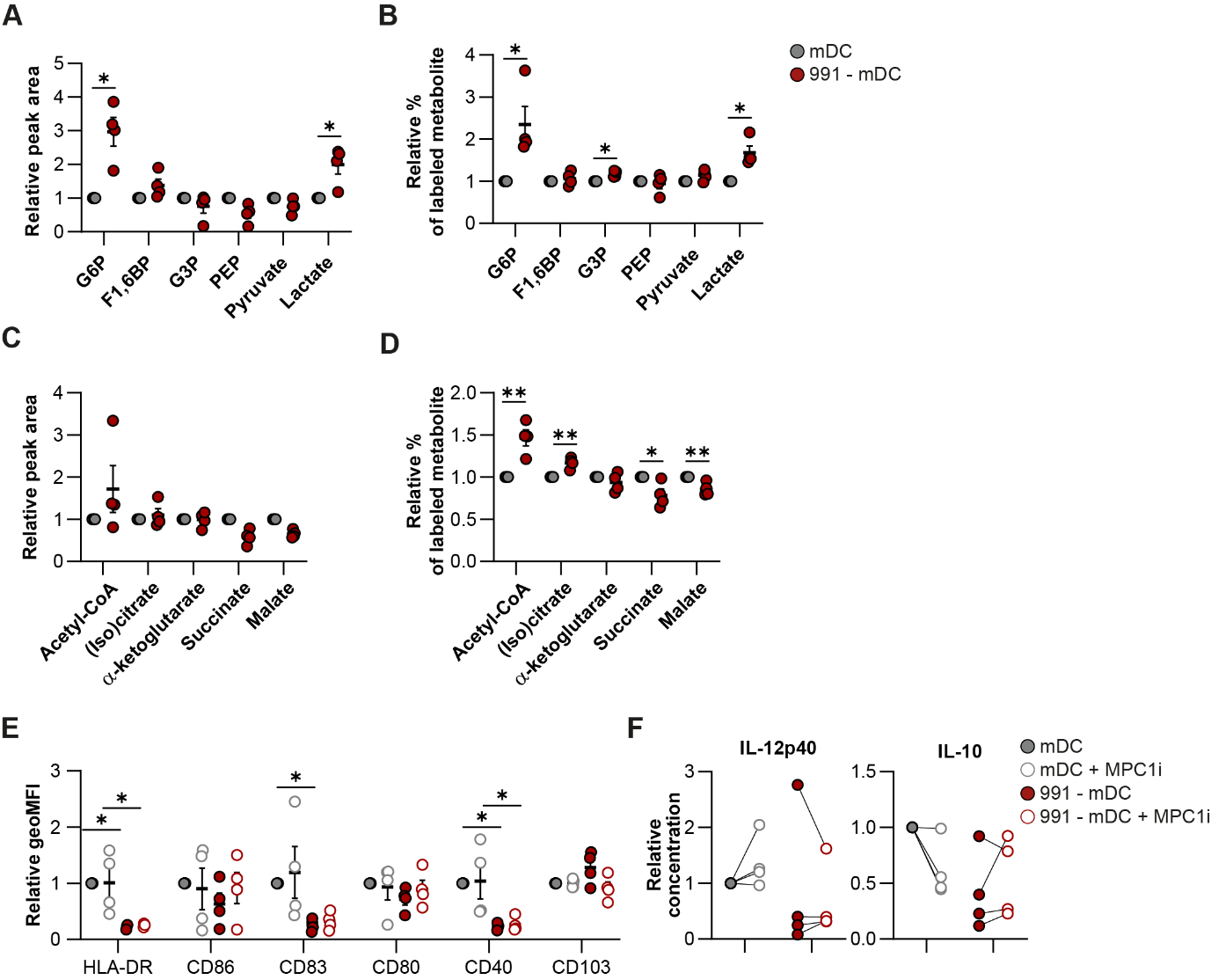

**Supplementary figure 5: effects of AMPK activation on glucose metabolism.**

**A-D:** Normalized abundance and normalized quantification of relative C13-glucose-labeled fraction of metabolites involved in **(A, B)** glycolysis and **(C, D)** TCA cycle, derived from figure 5A and B. **E:** Normalized expression of indicated markers. **F:** Normalized concentration of IL-12p40 and IL-10 after a 24hour co-culture with with CD40L-expressing J558 cells. Glucose 6-phosphate (G6P), fructose 1,6-biphosphate (F1,6BP), glycerol 3-phosphate (G3P), phosphoenolpyruvate (PEP). Results are expressed as means ± SEM. Datapoints represent independent experiments with different donors. Statistical analyses were performed using paired t-tests **(A-D)** or two-way Anova with Tukey post-hoc test **(E, F)**. *p < 0.05, **p < 0.01.
